# STING and VDAC inhibitors attenuate inflammation and ineffective erythropoiesis caused by an altered metabolome in the *Nan* (EKLF/E339D) mouse model of neonatal anemia

**DOI:** 10.64898/2025.12.11.693792

**Authors:** Tasleem Arif, Kaustav Mukherjee, Li Xue, Jonathan L Catrow, James Cox, James J Bieker

## Abstract

Erythroid Krüppel-like factor (EKLF/KLF1) is an essential transcriptional regulator of all aspects of erythropoiesis. The mouse neonatal anemia (*Nan*) mutation is driven by a semi-dominant mutation in one allele of the EKLF second Zn-finger at position E339D. RNA-seq analysis of Nan/+ erythroid cells showed that expression of numerous enzymes associated with metabolic pathways are changed as a result. We assessed and analyzed the effects of this dysregulation by mass spectrometry of embryonic and adult material. Our results show that mono-allelic expression of Nan-EKLF has profound impacts on erythroid cell metabolism: levels of amino acids, nucleotides, and metabolites are altered, more energy is needed for survival and proliferation, and glucose is taken up more rapidly. As a result, mitochondrial morphology is distorted, leading to VDAC1 oligomerization and mtDNA release to the cytosol. This activates the cGAS-STING signaling pathway and induces a type-I IFN response that drives inflammation. Use of STING or VDAC inhibitors alleviates these conditions both *ex vivo* and *in vivo*. Mechanistically, treatment restores normal erythroid cell divisions and differentiation, and decreases inflammatory pathways in the bone marrow. Our findings are likely directly relevant to the dyserythropoiesis observed in CDA type IV patients that carry a similar mutation.

## INTRODUCTION

Homeostasis of erythroid cells is a particularly critical task, where ∼2.5 million cells are produced per second in the human adult ^1^. Integration of intra- and extra-cellular signals is important for cell expansion and for the final, terminal differentiation steps of erythropoiesis, as these result in dramatic morphological changes that ultimately yield the mature, circulating erythrocyte ^1,2^. Coupled to this are fluxes in intracellular metabolites that can be readily changed, for example, in response to infection ^3^ or hypoxia ^4^. Critical for intracellular coordination and control of these events is Erythroid Krüppel-like Factor (EKLF; KLF1), a zinc finger transcription factor required for all aspects of erythropoiesis in both primitive and definitive erythroid cells ^5–11^. EKLF performs this function by binding its cognate 5’CCMCRCCCN recognition element via three carboxy-terminal C2H2 zinc fingers. EKLF expression is tissue-restricted throughout early development and in the adult, first appearing at the neural plate stage at E7.5, switching to the fetal liver by E9.5 and ultimately to the bone marrow and splenic red pulp of the adult ^12,13^. As a result, alteration of its function is predicted to affect erythropoiesis prenatally as well as in the adult. Ablation in the mouse leads to embryonic lethality by E14.5 due to a profound ß-thalassemia ^14,15^.

Links have been established between mutant KLF1 and altered human hematology and anemia ^5,16–18^, with mutations located throughout the gene ^8,19,20^. Many of those that lead to haploinsufficiency are benign but affect expression of a specific subset of its targets ^8,19,21–24^. However, a mutation at an orthologous amino acid in mice and humans leads to dominant anemia even when expressed as a heterozygote: congenital dyserythropoietic anemia (CDA) type IV in humans (E325K) ^25^, and neonatal anemia (*Nan*) in mice (E339D) ^26,27^. *Nan* is inherited in a semi-dominant fashion; homozygotes die in utero at E10-11, while heterozygous *Nan*/+ mice exhibit a lifelong severe anemia characterized by reticulocytosis, splenomegaly, altered globin expression, and cell membrane defects ^27–29^. This single amino acid change results in hypomorphic and neomorphic expression changes that alter red cell identity and its physiological properties, to the detriment of the organism ^30–32^. Molecularly, this is explained by the fact that the Nan-EKLF missense mutation is at a universally conserved amino acid in the second zinc finger, and that this change leads to altered DNA-binding properties *in vivo*. Two specific outcomes are that Nan-EKLF recognizes only a subset of WT sequences, and that it binds a new consensus binding sequence that is not normally associated with EKLF recognition, resulting in ectopic transcription. Mechanistically, Nan-EKLF interferes with WT EKLF binding, and RNA polymerase II pause-release properties are altered at both intrinsic and ectopic sites in Nan/+ erythroid cells ^33^.

An unexplained aspect is how the *Nan* mutation disrupts red cell metabolism, a particularly interesting question given its co-expression with WT protein and its dominant alteration of gene expression. Here, we evaluate the metabolomic consequences of cells expressing the Nan-EKLF mutation by examining erythroid cell metabolomes from E13.5 Nan/+ fetal liver erythroid cells, adult Nan/+ red blood cells and serum, and comparing to WT littermates. Our findings explain the Nan/+ red cell dominant anemia and abnormal erythroid phenotype, and likely model the dyserythropoiesis seen in human CDA type IV patients.

## RESULTS

### Altered gene expression related to metabolic pathways in the Nan/+ mutant fetal liver

Analysis of differential gene expression from bulk RNA sequencing between WT and Nan/+ FL erythroid cells ^30,32,33^ was used to evaluate global transcriptome changes related to alterations in the metabolome. These studies show that 195 genes have lower expression whereas 240 genes exhibit higher expression in Nan/+ ^33^. GO and panther pathway analyses show that the downregulated DEGs are mainly associated with cellular process, biological regulation, signaling, and developmental process, within which differentiation and development are highly enriched (Supplementary Fig. S1A). Interestingly, we also find that autophagy related genes are downregulated in Nan/+, which may lead to retention of dysfunctional mitochondria (addressed below).

At the same time, the upregulated, ectopic DEGs in Nan/+ compared to WT are mainly involved in metabolic processes, immune responses, and stress (Supplementary Fig. S1B). For example, the Ass1 gene (encoding argininosuccinate synthase) is one of the most highly upregulated genes in these cells (Supplementary Fig. S1C; ^32^), due to direct ectopic binding by Nan-EKLF to its promoter and intron, opening chromatin and enabling binding by CBP and H3K27ac ^33^. We have found that activation and repression of biological pathways by EKLF directly influence metabolic, inflammatory, and catabolic processes in Nan/+ FL cells.

### Identification of metabolic changes in Nan/+ FL cells by GC-MS

These dramatic changes in gene expression patterns compelled us to obtain a direct and comprehensive understanding of the metabolic changes in erythroid cells caused by the Nan/+ mutation. To better understand effects of changes in metabolism during erythroid development and to identify alterations in metabolites, we performed gas chromatography–mass spectrometry (GC-MS) analysis for comprehensive metabolic profiling of E13.5 FL cells (Fig. 1A). PCA plots show that FL cells from WT and Nan/+ are distributed in two separate areas, indicating a markedly different metabolome between them (Fig. 1B).

**Figure 1.**
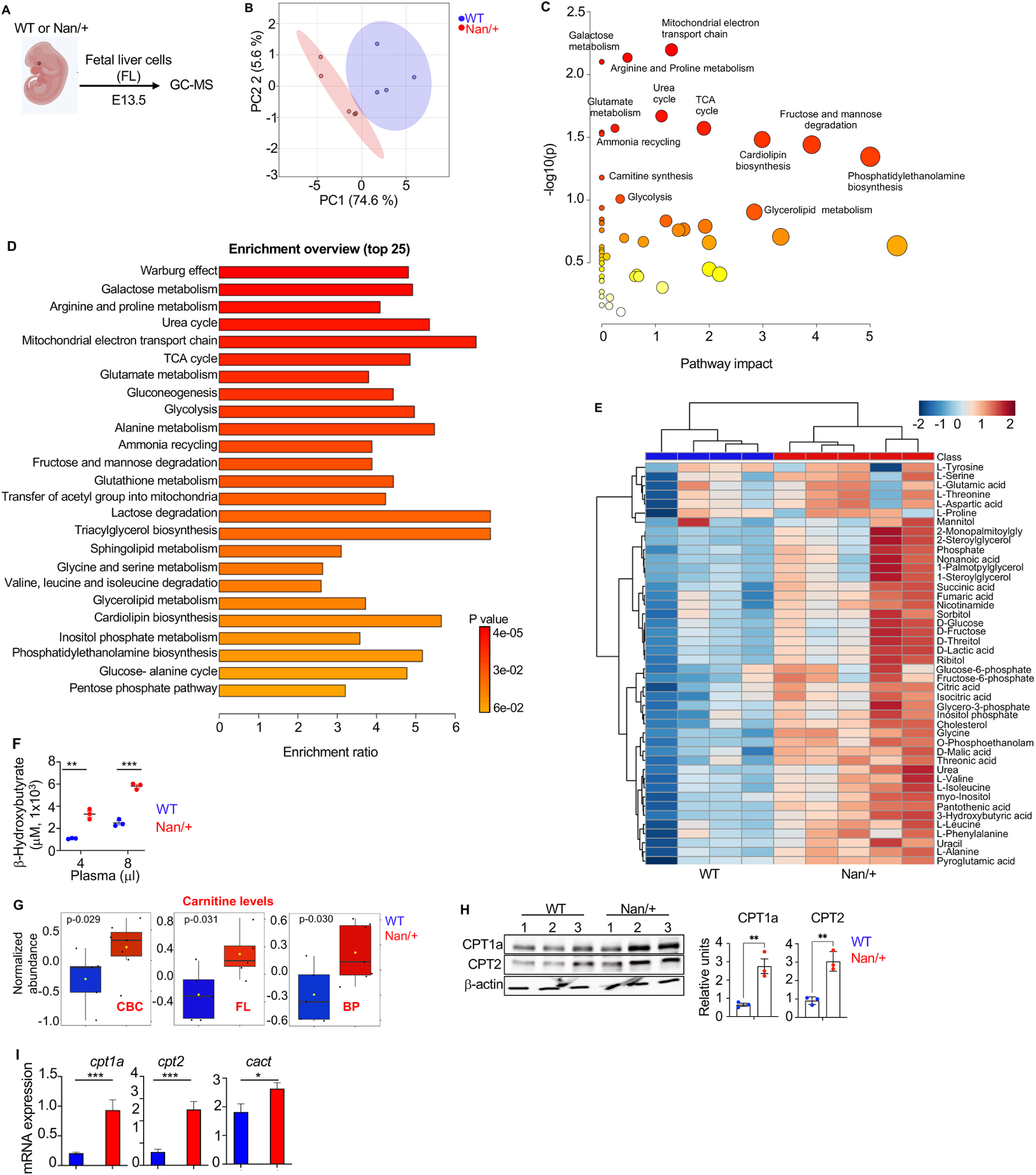
Global metabolic profiling of early developmental erythroid cells from WT and Nan/+ reveals significant differences in a wide range of metabolites (A) Study design for the metabolic investigation of erythroid cells. FL cells at embryonic day E13.5 from WT or Nan/+ mice were subjected to GC-MS analysis. (B) PCA plot shows analysis form WT or Nan/+ FL cells (WT in blue, Nan/+ in red), separating along PC1(74.6%) and PC2 (5.6%). Colored circles represent 95% confidence intervals and represent individual samples. (C) The pathway analysis module was used to generate the plot using the hypergeometric test and the latest KEGG version of the Mus musculus pathway library (MetaboAnalyst 5.0). Pathway topology analysis depict the most relevant pathways in the current set. The Y-axis defines the -log_10_ P-value, while the X-axis calculates the impact of the pathway in the metabolome analysis. The white to yellow to orange color gradation defines the significance of the pathway in the analysis, with red being the most significant in Nan/+ FL cells. The radius of the circle defines the number of metabolites obtained in the pathway. (D) An overrepresentation analysis (ORA) plot was generated in MetaboAnalyst 5.0. The plot shows the top 25 metabolic pathways that were enriched in Nan/+ FL cells. The ORA is the number of metabolic pathway metabolite hits divided by the expected number metabolites. Data are presented from n=6 to 7 biological replicates per group. (E) The heat map (distance measure using euclidean, and clustering algorithm using ward) depicts the top 40 metabolites obtained in analysis that are elevated in the WT or Nan/+ FL cells. **(F**). Concentration of β-OHB levels in BP of WT or Nan/+ adult mouse were analyzed by an ELISA assay. Data are expressed as the mean ± S.E.M. from n=4 biological replicates per group, **P < 0.01, ***P < 0.001. (G) Plot of normalized abundance concentration of L-carnitine levels significantly different metabolites (p < 0.05) in WT (blue bar) and Nan/+ (red bar) FL cells (**H**), CBC (**I**), and BP (**J)** obtained from WT or Nan/+. (H) Western blot analysis of the indicated proteins that play a role in carnitine shuttle pathway were performed using FL cell extracts from WT or Nan/+ (**left,** n=3) and their quantification (**right)**. β-actin was used as a loading/normalization control. (I) RT-qPCR analysis of indicated transcripts in freshly isolated FL cells from WT or Nan/+ (normalized to β-actin) (n = 3). (^∗^p < 0.05 and ****p < 0.001).

12 biochemical pathways are significantly altered in erythroid FL cells from Nan/+. The pathway impact plot shows that the most prominent are in the mitochondrial electron transport chain (ETC), TCA and urea cycles, and biosynthesis/degradation (Fig. 1C and Supplementary Table S5). The top 25 major pathways in FL cells identified in an MSEA graph are those belonging to mitochondrial metabolism (Fig. 1D and Supplementary Table S6). These results show that key energy producing metabolic pathways are dramatically altered in Nan/+ FL cells.

We more precisely identified modulated metabolites that are present exclusively in Nan/+ FL cells. A heatmap shows the top 40 metabolites found to be unique to Nan/+ (Fig. 1E, Supplementary Tables S7 and S8). Detailed analyses show that several metabolites related to mitochondrial metabolism, glycolysis and TCA cycle are exclusively enriched in Nan/+ FL cells (Fig.1E and Supplementary Tables S5-S8). The elevated level of glucose and lactic acid intermediate metabolites, together with the increase of TCA cycle, suggests rapid and high consumption of glucose during glycolysis in Nan/+ compared to WT FL cells and imply that mitochondrial activity is highly elevated in the Nan/+ FL cells.

We find several fatty acids (FAs) are significantly elevated in Nan/+ FL cells (Fig. 1E and Supplementary Tables S5-S8). The increase in FAs suggest Nan/+ FL cells are undergoing a higher level of mitochondrial β-oxidation than WT.

We further find that the level of the ketone body D-(−)-β-hydroxy butyric acid (also called β-HB) is increased in Nan/+ FL cells (Fig. 1E, and Supplementary Tables S5-S8), and that adult Nan/+ mice also have considerably increased circulation of β-hydroxybutyrate levels (Fig. 1F), suggesting a stable metabolic shift from glucose to fatty acid-derived ketones to provide enough energy for the survival of Nan/+ mutant mice.

### Metabolic profiling of Nan/+ mice reveals major changes in the circulating red blood cell and plasma metabolome

Liquid chromatography–mass spectrometry (LC–MS) and gas chromatography-mass spectrometry (GC-MS) were also used to identify distinguishing metabolites in circulatory red blood cells (CBC) and blood plasma (BP) sources from adult WT and Nan/+ mice to address whether the developmental metabolic changes suggested by the FL analysis (Fig. 1) persist even after birth.

Principal component analysis (PCA) following LC-MS shows significantly expressed metabolites identified in adult CBC (Supplementary Fig.S2A) and BP (Supplementary Fig.S2B). The results in Supplementary Fig S2C, 2D indicate that most of carnitine pathway, purine, pyrimidine, fatty acid, amino acid, and carbohydrate metabolites are elevated in Nan/+ CBC and BP samples (Supplementary Tables S1, S2, and S4). Consistent with the ectopic RNA observations, levels of argininosuccinate are significantly higher in Nan/+ CBC (Supplementary Fig. S2E).

In CBC, the concentration of several metabolites related to glycolysis and by-products of glycolysis pathways are significantly reduced in adult Nan/+ CBC (Supplementary Fig. S2F), whereas multiple metabolites related to the TCA cycle are enhanced (Supplementary Fig. S2G and Supplementary Tables S1-S2). Levels of several (but not all) amino acids (Supplementary Fig. S2H and Supplementary Tables S1-S2) and nucleotides (Fig. S2I and Supplementary Tables S1-S2), are significantly increased in Nan/+ CBC. Another metabolite, 4-aminobutyrate (GABA) that originates from glutamate and polyamines, is elevated in CBC of Nan/+ (Fig. S2J). Analysis of adult BP shows that other amino acids are significantly reduced in Nan/+ (Supplementary Fig. S2K and Supplementary Tables S3-S4), and multiple fatty acids are significantly elevated in adult Nan/+ BP (Supplementary Fig. S2L and Supplementary Tables S3-S4). These results indicate that metabolites related to TCA cycle and fatty acids, and selected nucleotides and amino acids are elevated, possibly due to genetic changes in Nan/+ mice or to altered uptake or stability.

The CBC analyses are also informative as related to energy requirements in the Nan/+ mice. We find pantothenic acid is elevated 24-fold in CBC of adult Nan/+ mice as compared to WT (Supplementary Fig. S2M, and Supplementary Tables S1-S2). This increase may be associated with higher transporting of acyl groups into the mitochondria for producing energy. The metabolite myo-inositol is also observed to be dramatically higher in Nan/+ adult mice (Supplementary Fig. S2M and Supplementary Tables S1-S2). The higher levels of myo-inositol suggest that Nan/+ mice are utilizing large amounts of glucose for the metabolism (addressed below). Finally, both CDP-choline and choline are significantly elevated in Nan/+ CBC as compared to WT (Supplementary Fig. S2N and Supplementary Tables S1-S2); these metabolites are degraded into uridine and play a critical role in acetylcholine and phospholipid metabolism ^34,35^.

These assessments, combined with the RNA expression analyses, demonstrate that the extensive genetic dysregulation seen within the Nan/+ erythroid cell leads to two significant global changes: first, it alters the erythroid metabolomic profile; second, these changes directly affect mitochondrial function. As a result, we focused the rest of our analyses on identifying the phenotypic consequences of these massive alterations of constituent metabolites on mitochondrial structure/function in Nan/+ cells.

### Carnitine levels and related genes are unexpectedly increased in Nan/+ mice

Particularly noteworthy are changes in carnitine levels in CBC, FL and BP (Fig. 1G) as well as individual carnitine variants (Supplementary Figs. S2C,D, S3A,B, and Supplementary Tables S1-S4), as verified by direct ELISA measurement of L-carnitine levels in BP (Supplementary Fig. S3C). Carnitine normally decreases during erythroid differentiation ^36^ as do other intermediate metabolites. L-carnitine is vital for the transport of fatty acids (FA) over the inner mitochondrial membrane (OMM) for FA β-oxidation, and production of ATP through the tricarboxylic acid (TCA) cycle ^37^.

To check whether genes involved in carnitine synthesis are altered in Nan/+ FL cells, we queried RNA samples from E13.5 FL as a suitable source for erythroid cells that are transcriptionally active, as this tissue is >90% erythroid ^38^. Carnitine shuttle pathway genes’ expression, such as CPT1a, CPT2, and CACT (Slc25a20), are increased in FL cells of Nan/+ at both protein (Fig. 1H) and transcript levels (Fig. 1I) levels. Similar results are seen by inspection of previously published bulk RNA sequencing data (not shown; ^32^), and these appear to be direct regulation targets ^30,33^. CPT1a is a rate-limiting enzyme that transports long-chain fatty acids into mitochondria for β-oxidation ^39^.

These metabolomic and DEG analyses suggest that the enhanced level and activity of CPT1A and/or CPT2 elevate the FA turnover rate in Nan/+ erythroid cells, implying that L-carnitine contributes to altered mitochondrial metabolism in Nan/+ mice in both the adult and early prenatal developmental stages (E13.5). As a result, we utilized the transcriptionally active and expanding E13.5 FL cells as our erythroid source for further in-depth analyses of global changes in metabolites and mitochondrial integrity.

### Mitochondria morphology and metabolic properties are altered in Nan mutant FL cells

Our hypothesis at this point was that the altered metabolite levels seen in Nan/+ lead to adaptive changes in the erythroid cell. To address this idea, we focused on three measurable properties of mitochondria: modification of morphology as related to metabolism, internal structure changes, and inflammatory responses that are predicted to follow such alterations.

First, given that energy-producing metabolic pathways are altered in Nan/+ FL cells as compared to WT, we assessed whether a variation in shape or size of mitochondria in this process might be a contributing factor to its cellular health. As expected, mitochondrial networks are fragmented in WT type FL cells (Supplementary Fig. 2A, S4A); however, mitochondrial fragmentation is low while surface area and volume are high in Nan/+ FL cells. This may be associated with decrease in autophagy related genes in Nan/+ FL cells (Supplementary Fig. S1A), which leads to retention of dysfunctional mitochondria. We measured mitochondrial mass after staining with MitoTracker Green (MTG), a probe that measures mass independently of mitochondrial membrane potential (MMP). We find mitochondrial mass is increased in Nan/+ FL cells compared to controls (Supplementary Fig 2B, S4B). Notably this increase in mitochondrial mass also leads to significant increases in mtDNA copy numbers (Fig. 2C and Supplementary Fig.S4C). These results suggest that mitochondrial activity is abnormally enhanced in Nan/+ FL cells.

**Figure 2.**
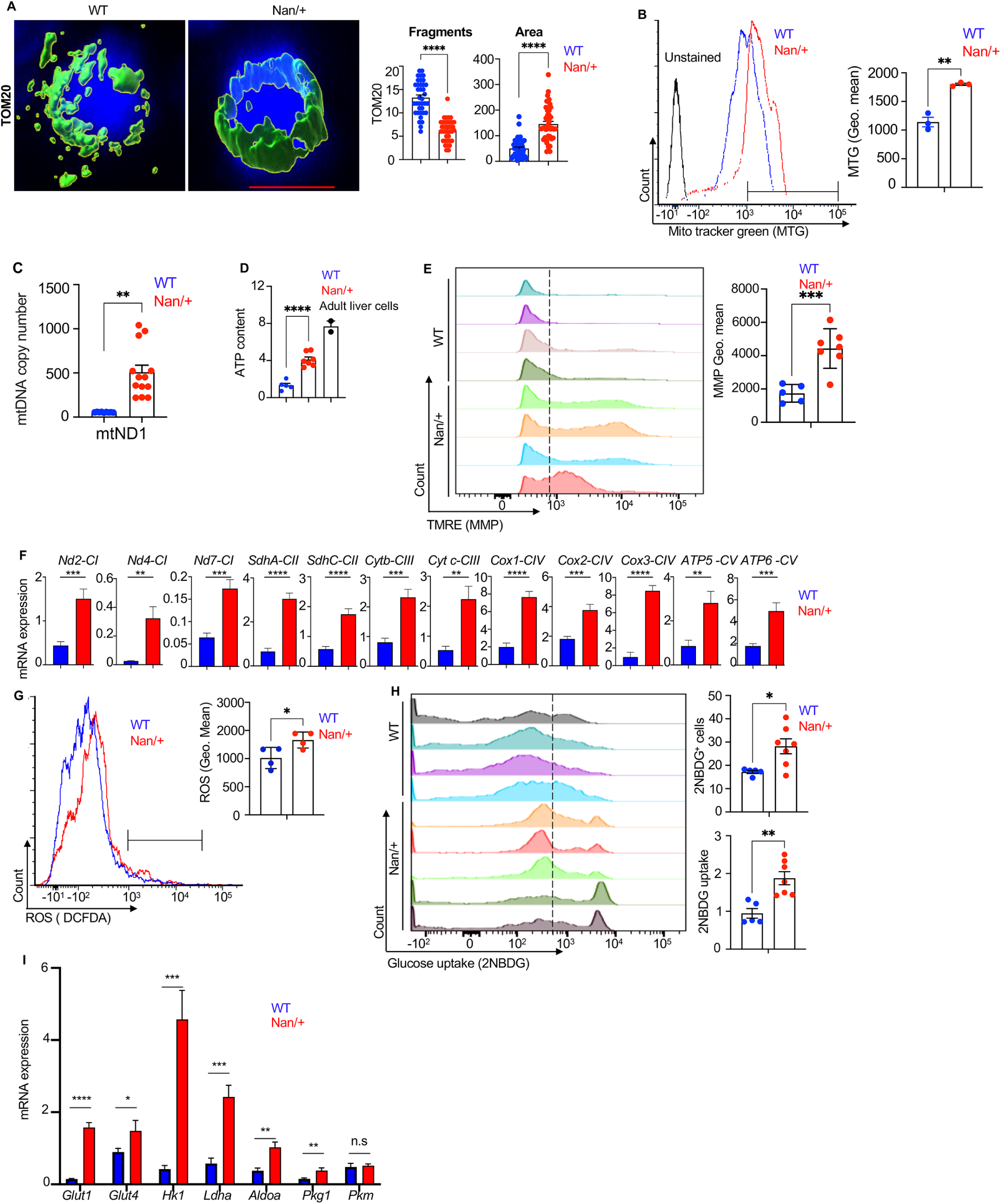
**Dynamic Regulation of Mitochondria in Nan/+ FL cells** Mitochondrial biogenesis and activities were analyzed in freshly isolated FL cells from WT or Nan/+. (A) Representative immunofluorescent confocal images of mitochondrial protein TOM20 showing the morphology of mitochondria (**left**, bar= 3 μm) and quantification of mitochondrial fragments and area **(right)**. (B) Representative FACS plot of mitochondrial mass measured in freshly isolated FL cells from WT or Nan/+ using MitoTracker Green (**left**), and quantification and the mitochondrial mass measured by the geometric mean (Geo.mean) fluorescence intensity (**right**). (C) Mitochondrial DNA copy numbers (mtDNA) was determined using RT-qPCR-based analysis to measure the ratio of mitochondrial DNA encoded protein NADH dehydrogenase 1 (mtND1) versus nuclear β-actin. (D) Mitochondrial ATP levels were measured in WT or Nan/+ FL cells (n = 4 mice). Gray bar represents cells isolated from adult liver that served as a control and for comparison. (E) Representative FACS plot (**left**) of changes in mitochondrial membrane potential (MMP) was analyzed by staining with TMRE. Quantification of MMP fluorescence levels is based on Geo.mean (**right**). (F) RT-qPCR analysis of electron transport chain (ETC) enzymes showing levels of indicated mRNAs encoding ETC transcripts of complex I, II, III, IV, and V (normalized to β-actin). (G) FACS analysis, and representative histograms of ROS levels (**left**) in FL cells from WT or Nan/+ were measured using CM-H2DCFDA dye. Quantification of ROS fluorescence levels based on Geo. mean (**right**). (H) Glucose uptake measured in freshly isolated FL cells from WT or Nan/+ treated with a glucose analog 2-[*N*-(7-nitrobenz-2-oxa-1,3-diazol-4-yl)amino]-2-deoxy-D-glucose, (2NBDG) for 2h in glucose-, pyruvate-, and glutamine- free media. Histograms (**left**) and quantification of percentage of 2NBDG positive cells (**right top**), and 2NBDG uptake are represented as mean fluorescence intensity (**right bottom**). (I) RT-qPCR analysis of metabolic enzymes showing levels of indicated mRNAs encoding glucose transporters and other glycolysis related transcripts (normalized to β-actin). Each experiment is an average of 3 to 7 biological triplicates per group. Data are presented as mean ± SEM (∗p < 0.05, ∗∗p < 0.01, ∗∗∗p < 0.001 and ∗∗∗∗p < 0.0001).

Mitochondrial dysfunction triggers an adaptive increase in mitochondrial biogenesis ^40,41^. Mitochondrial membrane potential (MMP) (ΔΨM) and cellular ATP generation were used to evaluate mitochondrial fitness. Mitochondrial respiration results in a high ΔΨM, whereas a decline in mitochondrial activity is linked to a decrease in ΔΨM. We find increased ATP levels (Fig. 2D) and increased MMP in Nan/+ FL cells (Fig. 2E and Supplementary Fig. S4D) that indicate the cell’s attempt to restore normal mitochondrial function that has been compromised in Nan/+ FL cells.

The decrease in mitochondrial fragmentation and enhanced mitochondrial activity in Nan/+ FL cells with elevated mitochondrial metabolism may be due to the elevated expression of mitochondrial-specific genes, including those of the electron transport chain (ETC). qRT-PCR analysis confirms that the expression of all ETC complexes’ genes is significantly higher in Nan/+ FL cells (Fig. 2F). Notably, respiratory chain complexes I and III are the primary sources of mitochondrial reactive oxygen species (ROS) ^42,43^. We find significantly higher levels of cytosolic ROS in Nan/+ FL cells (Fig. 2G and Supplementary Fig. S4E). Enhanced ROS levels are reported to be associated with several blood disorders including hemolytic anemia ^44,45^, a phenotype shown by Nan/+ mice and CDA IV patients.

Since alteration of mitochondrial morphology influences glucose metabolism ^46^, we next asked whether this impacts glucose consumption in Nan/+ FL cells. Glucose uptake, measured using 2NBDG under metabolic (pyruvate, glucose and glutamine)-free conditions ^47^, is elevated 2-fold higher in Nan/+ when compared to WT FL cells (Fig. 2H and Supplementary Fig. S4F). qRT-PCR analysis further confirms that the expression of glycolysis-related genes, including Glut1 (Slc2a1) and Glut4 (Slc2a4), and the main glucose transporter Hk1, are all expressed greater in Nan/+ FL cells (Fig. 2I), possibly via a direct regulation ^30,33^. Other molecules involved in glucose metabolism are also increased such as Ldha, Aldoa, and Prkg1 (Fig. 2I; however, Pkm is not changed). These data suggest a means by which glucose might be transported at a higher level in Nan/+ erythroid cells.

Overall, these results support the notion that Nan/+ FL cells have elevated mitochondrial biogenesis that follows from altered fragmentation, membrane potential, and gene expression, resulting in use of more energy from increased glucose metabolism for their survival and longevity.

### Cristae disorganization induces mtDNA release and induces mtDNA-dependent IFN-I response in Nan/+ mutants

The morphology of mitochondria is dynamic, changing both at the cellular and intra-mitochondrial levels in response to energy and/or developmental cues ^48,49^. The second part of our hypothesis was to assess to what extent mitochondrial internal phenotypic structures are altered in Nan/+. To address this, we examined mitochondrial morphology by transmission electron microscopy (EM). Unexpectedly, we find that Nan/+ FL cells exhibit enhanced, but variable, degrees of cristae disorganization, such as loss of cristae, detachment of inner membrane from outer membrane, and mitochondrial vacuolization (Fig. 3A and Supplementary Fig. S5A) in contrast to WT FL cells. The morphology of mitochondrial cristae helps protect against mtDNA release and inhibits mtDNA-dependent IFN-I response to induce inflammation ^50^. Irf7 is a transcription factor primarily expressed by macrophages that mediates inflammation by activating interferon β (IFN-β) ^51,52^. Irf7 expression in the Nan/+ FL, spleen, and bone marrow erythroid cells is higher than WT, resulting in increased levels of serum IFN-ß, thus repressing erythropoiesis ^32^. Consistent with this, Panther gene ontology (GO) analysis of published RNA-seq data ^30,32^ reveals that biological processes such as inflammatory and immune system responses are significantly enriched in upregulated genes of Nan/+ FL cells (Supplementary Fig. S1B). Furthermore, expression and release of pro-inflammatory cytokines can be augmented by the CPT1A overexpression or increased activity we have observed in Nan/+ FL cells (Supplementary Fig. S2I,J).

**Figure 3.**
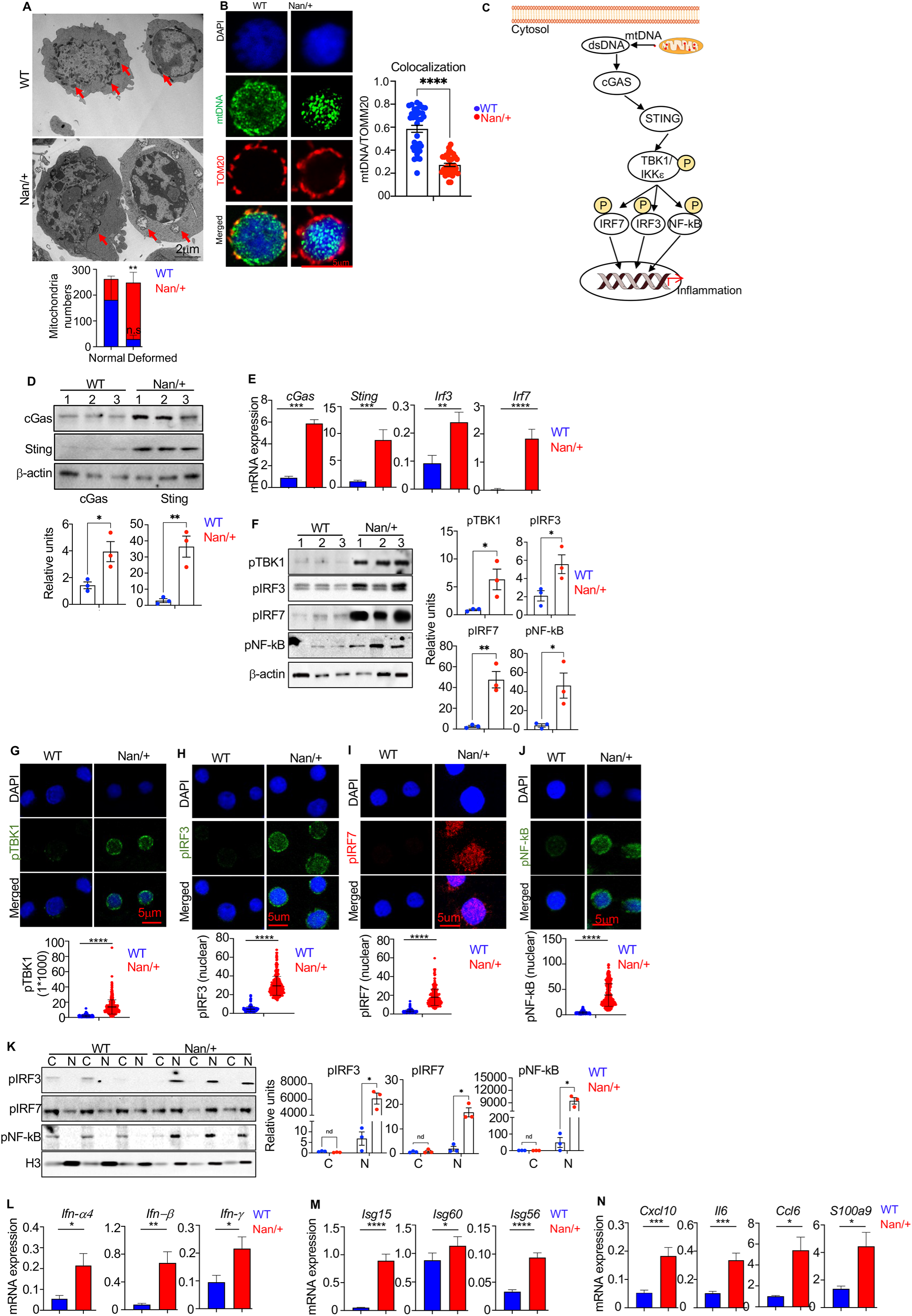
**Deformed mitochondrial cristae and STING signaling pathway is activated in Nan/+ FL cells** (A) Representative transmission electron microscopy (EM, **top**) images of mitochondria in FL cells from WT or Nan/+ reveals normal-shaped and -size mitochondria in WT mice while Nan/+ mice show enhanced, but variable, degrees of cristae disorganization, such as cristae loss, detachment of inner membrane from outer membrane, and mitochondrial vacuolization. Quantification of deformed mitochondria (**bottom**) (bar = 2 µm). (B) Representative immunofluorescent super resolution confocal images of mtDNA colocalization with mitochondria protein TOM20 in freshly isolated FL cells from WT or Nan/+ (**left**). Quantification of mtDNA and TOM20 colocalization, (**right**; bar= 5μm). (C) A schematic detailing mtDNA-induced activation of the mtDNA-GAS-STING pathway. STING recruits TANK-binding kinase 1 (TBK1), promoting TBK1 autophosphorylation, and recruitment of interferon regulatory factor-3 and-7 (IRF-3, and -7). The phosphorylation of IRF-3 or -7 by TBK1 enables IRF-3, -7 and NF-kB p65 protein translocation to the nucleus to induce gene expression of type I interferons, thus inducing inflammation. (D) Western blots of whole cell extracts showing levels of cGAS-STING pathway proteins in freshly isolated FL cells from WT or Nan/+ (**top** ), and quantification as relative units to β-actin (**bottom**). (E) RT-qPCR expression analysis of indicated STING pathway transcripts (normalized to β-actin). (**F**) Western blots of whole cell extracts showing levels of STING pathway proteins in freshly isolated FL cells from WT and Nan/+ (**left**). Quantitative analysis presented as RUs normalized to β-actin (**right**). (**G-J**) Representative confocal images showing immunofluorescent analysis of STING pathway related proteins (bar = 5μm: pTBK1 (**G, top**); nuclear translocation of pIRF3 (**H, top**); pIRF7 (**I, top panel**); and pNF-kB65 (**J, top**) in freshly isolated FL cells from WT or Nan/+ and their quantification (**G-J**; **bottom**). (**K**) Nuclear and cytoplasmic fractions were isolated from fresh FL cells from WT or Nan/+, and were analyzed by immunoblot for pIRF-3, -7 and pNF-kB p65 localization (l**eft)**. Fractionation efficiency was quantified by localization of H3. Quantitative analysis is presented as relative units normalized to H3 (**right**). (**L-N**) RT-qPCR analysis of expression of indicated type I interferon (**L**), interferon stimulatory genes **(M),** cytokines and chemokines (**N**) in freshly isolated FL cells from WT or Nan/+ (normalized to β-actin). Each experiment is an average of 3 to 5 biological triplicates per group. Data are presented as mean ± S.E.M. (^∗^p < 0.05, ^∗∗^p < 0.01, and ^∗∗∗^p < 0.001, and ****p < 0.0001).

Alternatively, elevated inflammatory gene expression can also be triggered by mitochondrial dysfunction ^53^, fostering the notion that mitochondrial damage and enhanced mitochondrial metabolism might also contribute to the elevated expression of these inflammatory pathways ^54^. The third part of our hypothesis suggests that such damaged mitochondria in Nan/+ might trigger inflammation via activation of the mtDNA-cGAS-STING pathway. Given this possibility in addition to the elongated mitochondrial network and deformed mitochondrial cristae (Fig. 2A, 3A and Supplementary Fig. S4A, S5A), we tested whether damaged mitochondria and removal of mitochondrial DNA (mtDNA) may be compromised and thus detected in Nan/+ cytosol, outside of mitochondria. Analysis of data points from super resolution confocal immunofluorescence (IF) show that in Nan/+ FL cells, mtDNA escapes mitochondria, and its mitochondrial localization is significantly reduced compared to WT (Fig. 3B and Supplementary Fig. S5B). Moreover, several molecules involved in the mtDNA-cGAS-STING pathway (Fig. 3C) are significantly elevated at protein (Fig. 3D,F) and transcript (Fig. 3E) levels in Nan/+ FL cells. Leakage of mtDNA and ROS production could synergistically contribute to the activation of mtDNA-cGAS-STING pathway ^55^. Together these findings suggest that the mtDNA-cGAS-STING signaling pathway is activated in Nan/+ FL cells.

We further probed the potential engagement of the cGAS-STING signaling pathway by mtDNA stress and find that while STING recruitment and phosphorylation of its substrate TBK1 is almost non-existent in WT, pTBK1 is greatly elevated in Nan/+ FL cells (Fig. 3G and Supplementary Fig. S5C). Furthermore, the phosphorylated forms of TBK1 downstream targets, IRF3 and IRF7, are prominently elevated and accumulate in the nucleus of Nan/+ FL cells (Fig. 3H,I), indicating their activation, whereas pIRF3 and pIRF7 are mainly absent or less expressed from the nucleus in WT FL cells (Fig. 3H,I and Supplementary Fig. S5D,E).

The inflammatory response is a central pathological process, and a key mediator (including in the erythroid cell) is nuclear factor κB p65 (NF-κB-p65), another TBK1 substrate ^56^. Many inflammation triggers, including released mtDNA, are known to activate NF-κBp65 and many effectors of inflammation such as inflammatory cytokines are activated either directly or indirectly by NF-κBp65. Immunostaining of Nan/+ FL cells reveals a dramatic increase in levels of phosphorylated NF-κBp65 and its localization in the nucleus of Nan/+ cells compared to WT (Fig. 3J and Supplementary Fig. S5F). Separation of cell extracts into nuclear/cytosolic fractions and analysis by western blot show that p65 protein is enhanced in the nucleus of Nan/+ FL cells along with pIRF-7 and -3 proteins (Fig. 3K). Activation of this factor thereby increases transcripts for IFNs (IFN-α4, IFN-ß and IFN-ψ) (Fig. 3L), interferon response genes (ISG15, ISG60 and ISG56) (Fig. 3M), and several inflammatory chemokine and cytokines (Fig. 3N) in Nan/+ FL cells.

Together these findings show that the mtDNA-cGAS-STING signaling pathway is activated in Nan/+ mutant FL cells because of disrupted mitochondrial cristae structure and mtDNA leakage, leading to an inflammatory response.

### STING inhibitor reduces expression of inflammatory genes in culture and improves anemia of Nan/+ mice in vivo

To evaluate the contribution of inflammation to the anemia, we treated in vitro cultures of Nan/+ or WT FL (E13.5) cells with pharmacological STING inhibitors (H151 or C176) and assayed erythroid maturation (Fig. 4A, left). We find that compared to WT, RNA expression of IFNs (-α4, -ß, and -ψ) (Fig. 4A, bottom), ISGs and inflammatory genes (Supplementary Fig. S6A) are reduced several fold specifically in Nan/+ FL cells treated with the STING inhibitors. Key enzymes involved in activation of the cGAS-STING pathway, such as pTBK-1, p-IRFs (-3, -7) and NF-κB-p65 are also repressed in Nan/+ upon H151 treatment (Fig. 4A, right). Ex vivo H151 treatment also reduces the number of early-stage erythroblasts (CD44-hi Ter119-) and increases late-stage/mature erythroblasts cells (CD44-lo Ter119+) (Fig. 4B and Supplementary Fig. S6B). We find maturing erythroid populations in Nan/+ only upon H151 treatment (Supplementary Fig. S6C), whereas these populations are low/absent in DMSO control-treated Nan/+ cells in time dependent manner.

**Figure 4.**
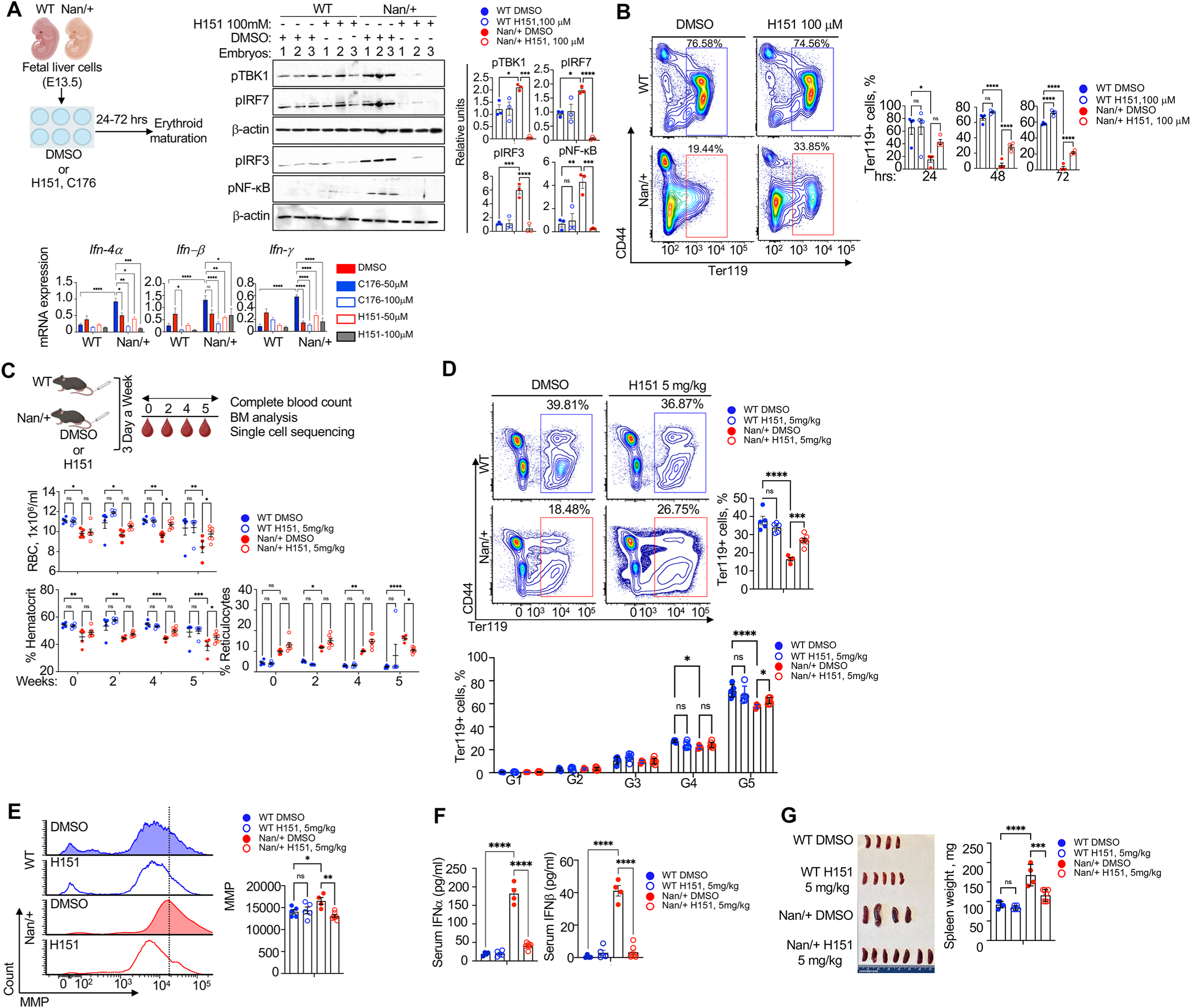
**STING inhibitor ameliorates anemia in adult Nan/+ mice** (A) Schematic diagram showing sting inhibitor (H151 and C176) treatment workflow in FL (E13.5) derived from WT or Nan/+ (n=3). FL cells were incubated in the presence of DMSO (control) or STING inhibitors (50 to 100μM) for 24-72 hours. Western blots of extracts probed with anti- pTBK1, -pIRF7, -pIRF3 and pNF-kB-p65 for biological triplicates are shown after H151(100μM) treatment and their quantification is shown (right); β-actin served as a loading control (middle). Relative mRNA levels of Ifn-4α, -ß, or -ψ from samples treated with two concentrations of STING inhibitors H151 (50-100 μM), and C176 (50-100 μM) are as indicated (bottom) (B) Representative flow plot of FL cells treated with DMSO or STING inhibitor ex vivo were analyzed for erythroid progression by CD44/Ter119 at different times (24 to 72 hrs, left) and quantification of Ter119+ percent (right). (C) Schematic diagram showing STING inhibitor (H15, 5 mg/kg) treatment workflow. Mice ( n= 5 to 6) were injected IV with DMSO (control) or H151 and blood samples analyzed at every two weeks of interval (top), followed by total harvest of bone marrow at the end for blood parameters such as RBC numbers (top middle), hematocrit ( left bottom) reticulocyte counts (right bottom), and single cell analyses (of Figs 5 and 6). (D) Representative flow plot of bone marrow cells from DMSO or H151-treated mice were analyzed for Ter119+ (top) as well as well as erythroid maturation using CD44/FSC (forward scatter) of Ter119+ cells and gating them into groups G1-G5 based on high to low CD44/FSC as indicated (bottom, gating strategy in Figure S6C). (E) MMP analyzed in BM Ter119 + cells treated with DMSO or H141; histograms (left) and quantification (right). (F) Serum IFN-α (left) and -ß (right) levels measured using Elisa in mice treated with DMSO or H151. (G) Spleens were isolated and weighed. Size is shown (**left**) and weight is quantified (**right**). Each experiment is an average for ex vivo (n=3) and in vivo (n=5 to 6). In all panels, data are presented as mean ± S.E.M. (^∗^p < 0.05, ^∗∗^p < 0.01, and ^∗∗∗^p < 0.001, and ****p < 0.0001).

Next, *in vivo* effects of STING inhibition were assessed by intravenous treatment of adult mice with H151 followed by removal of blood at various time points (weeks) (Fig. 4C). STING inhibitor treatment significantly alleviates anemic hallmarks as seen by increased RBC count and hematocrit, and decreased reticulocyte counts in Nan/+ mice, particularly after 5 weeks (Fig. 4C, bottom). H151 treatment improves ineffective erythropoiesis in the bone marrow (BM) of Nan/+ mice by enhancing the number of Ter119+ positive cells (Fig. 4D) and mature erythroid cells (Fig. 4D, and Supplementary Fig. S6D). Several key enzymes involved in activation of the cGAS-STING pathway, such as pTBK-1, p-IRFs (-3, - 7) and NF-kB-p65 are reduced in BM Ter119+ cells (Supplementary Fig S6E), and such treatment further reduces and normalizes the high mitochondrial activity seen in Nan/+ FL cells (Fig. 4E). Consistent with our transcript data we find IFNα and INFß protein levels in serum are reduced in Nan/+ treated with H151 compared to WT (Fig. 4F). Finally, splenomegaly is reduced in the Nan/+ mice after H151 treatment (Fig. 4G). Together these data show that the Nan/+ anemia phenotype can be alleviated in culture and in vivo by using GAS-STING inhibitors to decrease inflammatory pathways.

### H151 treatment enhances normal cell cycle specifically in the in vivo erythroid population and restores effective erythropoiesis in Nan/+ bone marrow (BM)

To examine the effects of H151 treatment on BM hematopoiesis in WT and Nan/+ adult mice, we performed single cell mRNA sequencing on pooled total BM cells from 3 mice of each genotype using the 10X Chromium platform. Hematopoietic populations were resolved using PCA and UMAP projection, and clusters were labeled based on marker gene expression ^57^;Stumpf, 2020 #3126} (Supplementary Fig. S7A). Non-erythroid populations were unaffected upon treatment with H151 in both WT and Nan/+ (Supplementary Fig. S7B), whereas a subset of the early erythroid population showed an altered clustering pattern in the Nan/+ BM treated with H151 (Supplementary Fig. S7B, black circle). Next, the early and late erythroid populations were re-clustered (Fig. 5A) and erythroid stages were determined based on gene expression (Supplementary Fig. S7C). Upon visualizing the populations separately by sample and condition, we find that untreated Nan/+ BM shows a different profile than the other samples likely due to altered gene expression, and orthochromatic erythroblasts are drastically reduced in Nan/+ (Fig. 5A). Upon H151 treatment, most of the cells in Nan/+ revert to normal WT-like populations whereas WT is unaffected by H151 treatment (Fig. 5A) indicating that H151 specifically affects the erythroid cells in Nan/+.

**Figure 5.**
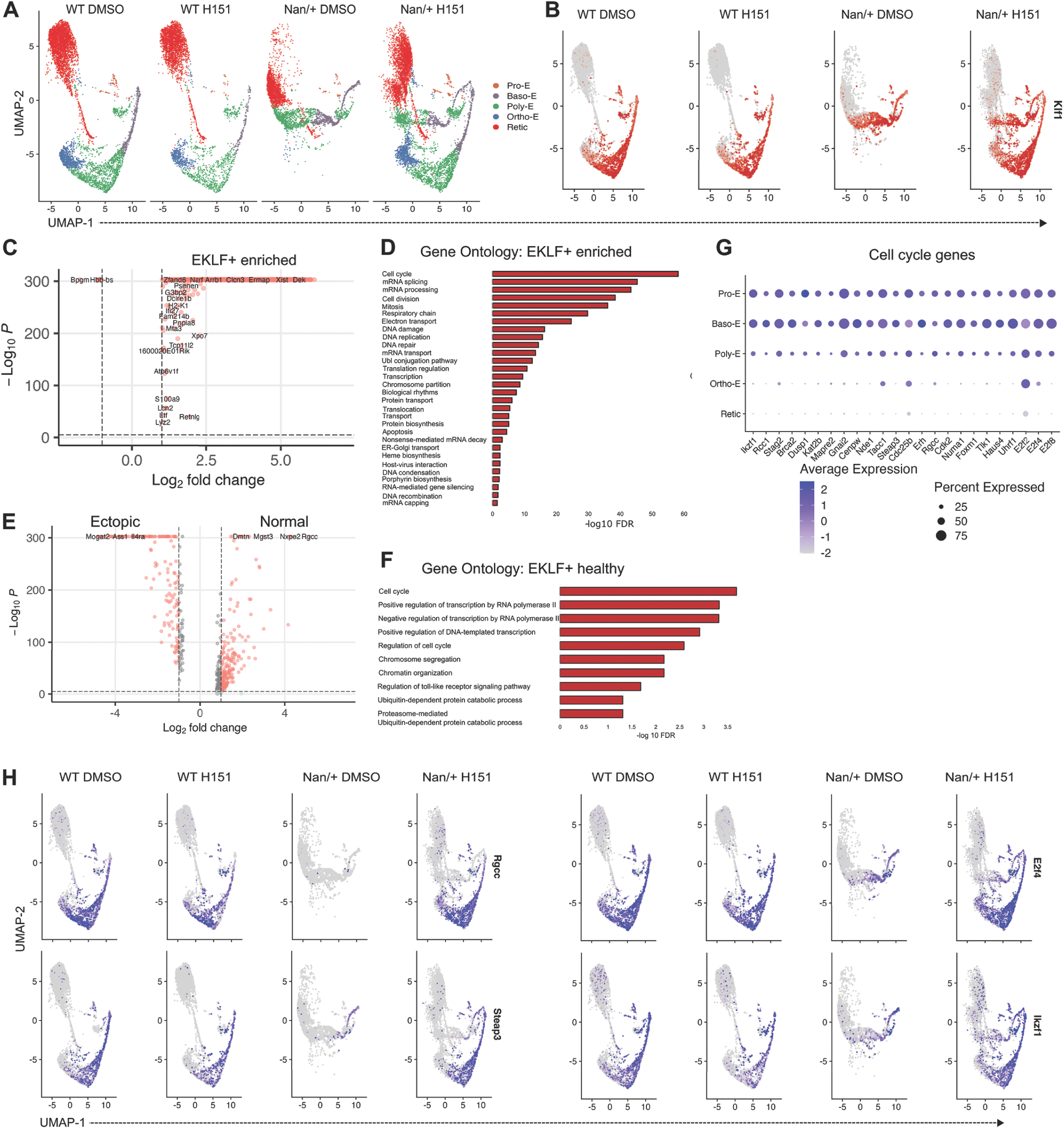
Single cell analysis of WT and Nan/+ bone marrow after treatment with the STING inhibitor H151. **(A)** Clustering and U-MAP projection of early and late erythroid sub-populations from Figure S7A separated by genotype and H151 treatment. Clusters are colored by cell-type indicated in the legend. **(B)** U-MAP projection of erythroid clusters separated by sample showing EKLF/Klf1 expressing cells as red dots. Intensity of red color indicates expression level of EKLF. **(C)** Volcano plot showing the differentially expressed genes in the EKLF+ and EKLF- erythroid cells with specific genes labeled. **(D)** Gene ontology analysis of genes with enriched expression in the EKLF+ erythroid populations. **(E)** Volcano plot showing differentially expressed genes in the healthy WT-like EKLF+ erythroid populations and Nan-like EKLF+ populations with ectopic gene expression. **(F)** Gene ontology analysis of genes with enriched expression in the healthy EKLF+ erythroid populations. **(G)** Dot plot showing the expression level and percentage of cells expressing indicated cell cycle genes in the various erythroid populations in all the samples. **(H)** Feature plot showing the expression of specific cell cycle EKLF target genes in the individual samples.

Next, we identified the EKLF-expressing population (Fig. 5B) and performed pseudobulk differential expression analysis to compare EKLF+ and EKLF- cells (Fig. 5C). We find that genes enriched in EKLF+ clusters (Supplementary Table S9) are mostly involved in cell cycle and other cellular functions that support the rapid cell proliferation required during erythroid maturation (Fig. 5D), whereas the EKLF-clusters are enriched for markers of mature erythroid cells (Supplementary Table S10). Next, we analyzed the EKLF+ cells in Nan/+ versus WT (independent of H151 treatment) (Fig. 5E) which reveals that differential gene expression in BM largely resembles our earlier bulk RNA-seq analysis of WT and Nan/+ fetal liver (Supplementary Fig. S7D ^32^). The healthy EKLF+ cells of WT mostly express genes (Supplementary Table S11) involved in cell cycle and cell division and associated molecular processes (Fig. 5F). In contrast, the unhealthy Nan-like EKLF+ cells express genes involved in cytoskeleton organization and apoptosis indicating that many EKLF+ cells in Nan/+ BM maybe apoptotic leading to persistent anemia in adulthood (Supplementary Table S12). Expression of cell cycle genes correlates with EKLF expression in the populations (Figs. 5G, 5B and Supplementary Fig. S7C) and some EKLF target cell cycle genes lose expression in Nan/+ but strikingly, their expression is restored upon H151 treatment of Nan/+ BM (Fig. 5H). We performed a cell cycle scoring on the populations in WT and Nan/+ and found that cell cycle stages were identical in the two samples indicating that observed clustering differences were not due to cell cycle stage (Supplementary Fig. S7E). The above observations indicate that H151 treatment prevents apoptosis of erythroid cells in the Nan/+ BM and restores normal erythroid cell division. The anemic environment in Nan/+ BM is driven by ectopic gene expression in erythroid cells leading them to cluster separately from the healthy cells in WT (Supplementary Fig. S7F). Since H151 inhibits cGAS-STING driven expression of Type I inflammation genes, examination of genes like Irf2, Irf3, Irf7, and Cdkn1a show that unhealthy Nan-like erythroid cells expressing these genes are reduced upon H151 treatment and restored to a healthy WT-like profile (Supplementary Fig. S7G).

Next, we examined differential gene expression of unhealthy erythroid cells in the Nan/+ BM with and without H151 treatment to find that certain genes have higher expression after H151 treatment (Fig. 6A, Supplementary Table S13). These genes are involved in autophagy, immunity and apoptosis (Fig. 6B), which suggests that H151 treatment may also accelerate apoptosis and autophagy of unhealthy erythroid cells. Further, reclustering of erythroid clusters in Nan/+ with and without H151 treatment (Fig. 6C) and erythroid marker expression analysis (Fig. 6D) reveals that markers associated with mature erythroid cells are present only in the untreated Nan/+ ectopic subset as depicted by clusters 0, 2, and 7 (Fig. 6E, left). This suggests that mature unhealthy erythroid cells are absent after H151 treatment of Nan/+ BM. Separating by sample also reveals two mutually exclusive cell populations – Cluster 4 in Nan/+ and Cluster 1 in H151 treated Nan/+ both of which are EKLF+ (Fig. 6E, right). Differential gene expression in these clusters (Fig. 6F) reveals that genes enriched in cluster 4 of Nan/+ are cell cycle genes (Fig. 6G, Supplementary Table S14) indicating that these are actively dividing cells, but their reduced numbers compared to WT or Nan/+ treated with H151 explains the low number of mature erythroid cells in the Nan/+ BM. In contrast, the cells in cluster 1, present only upon H151 treatment, are enriched for immune response and inflammatory genes (Fig. 6H, Supplementary Table S15), many of which are expressed in earlier stages of myeloid/erythroid progenitors ^58^.

**Figure 6.**
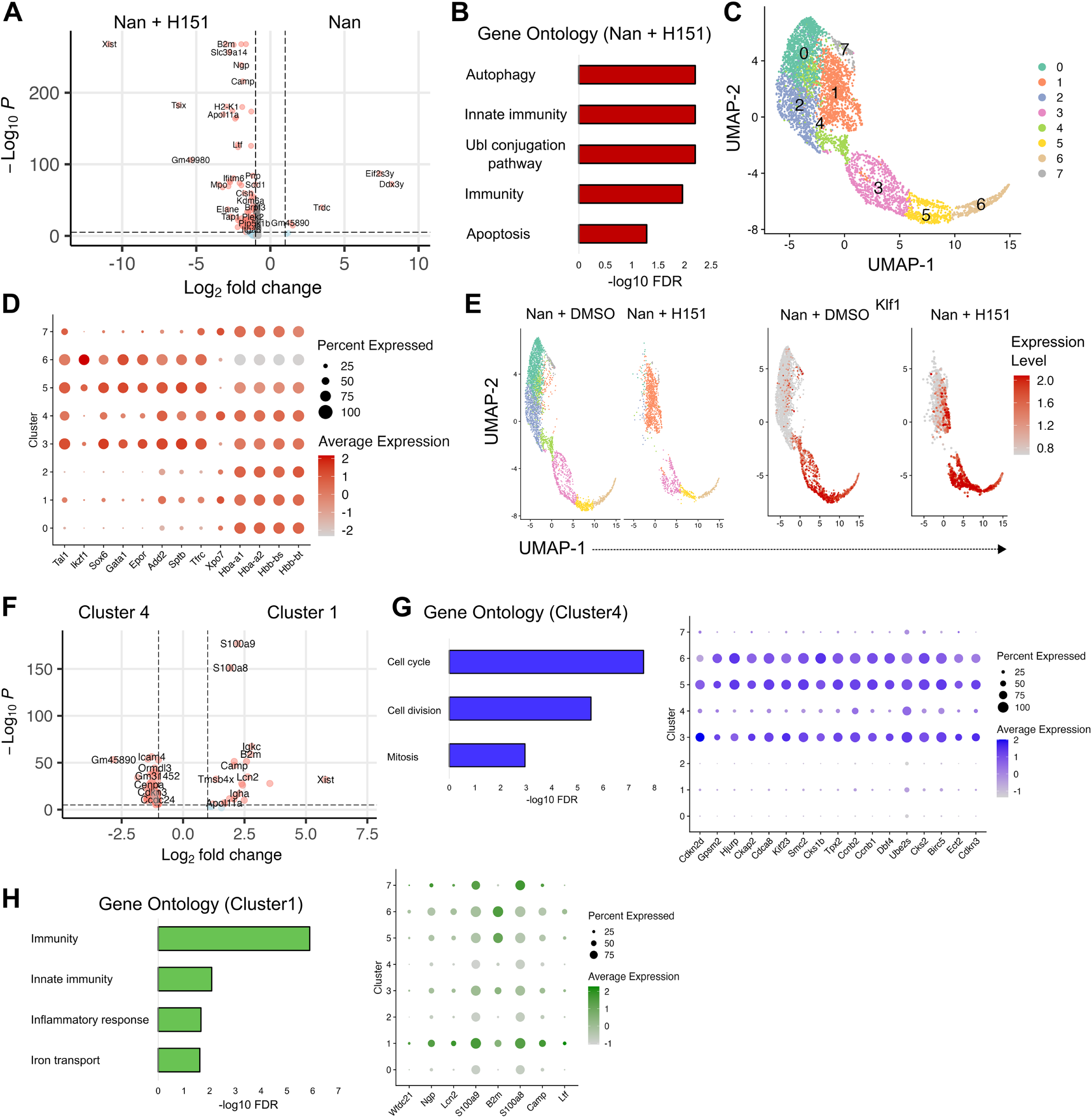
Single cell analysis of unhealthy erythroid cells with ectopic gene expression in Nan/+ bone marrow. **(A)** Volcano plot showing differential gene expression in the erythroid cells of the Nan/+ bone marrow with and without H151 treatment. **(B)** Gene ontology categories of genes with enriched expression in unhealthy erythroid cells of Nan/+ bone marrow after H151 treatment. **(C)** Reclustering and U-MAP projection of the unhealthy erythroid cells in Nan/+ with and without H151 treatment. **(D)** Dotplot showing expression of specific early and late erythroid genes in the various clusters indicated in (C). **(E)** U-MAP projection of clusters from (C) separated by sample (left) and showing EKLF expression as indicated in the legend (right). **(F)** Volcano plot showing differentially expressed genes in clusters 4 and 1 identified in (C). **(G)** Gene ontology analysis showing biological functions of genes enriched in cluster 4 (left) and dotplot showing specific cell cycle genes enriched in cluster 4 (right) . **(H)** Gene ontology analysis showing biological functions of genes enriched in cluster 1 (left) and dotplot showing specific immune response genes enriched in cluster 1 (right).

These results indicate that the erythroid cells in Nan/+ BM continue to undergo cell division and erythroid maturation despite ectopic gene expression, which leads to unhealthy maturation without transitioning through an orthochromatic erythroblast stage. H151 treatment inhibits inflammatory pathways specifically in the Nan/+ BM erythroid population, reduces the number of unhealthy erythroid cells through apoptosis, and restores normal erythroid cell divisions and populations to rescue adult Nan/+ mice from persistent anemia.

### Elevated mitochondrial stress and mitochondrial metabolism lead to the release of mitochondrial DNA

We have shown that mitochondrial stress, triggered by increased mitochondrial metabolism following deformed mitochondrial cristae (Fig. 2A and Supplementary Fig. S5A) ^50^, leads to the release of mitochondrial DNA (mtDNA) into the cytosol in Nan/+ FL cells (Fig. 3B and Supplementary Fig. S5B). Subsequently, mtDNA interacts with and activates a large number of immunostimulatory DNA sensors, such as cyclic GMP-AMP synthase (cGAS), that can trigger a type-I interferon (IFN) response ^59^. As a result, we addressed the mechanism by which these observations are linked in the Nan/+ mitochondria.

It is known that mitochondrial outer membrane permeabilization (MOMP) is needed for the release of mtDNA into the cytosol ^60^. BAX/BAK oligomers, which can form macropores in the mitochondrial outer membrane (MOM), also facilitate mtDNA release ^61–63^. However, BAX/BAK macropore formation is usually formed under conditions that activate BAX/BAK, such as apoptosis or treatment with their activators ^61–63^. The pore(s) that stimulates MOMP in live cells or in circumstances that do not engage BAX/BAK have not been determined. We examined whether Nan/+ FL cells are undergoing apoptosis at a steady state. Annexin V/PI staining indicates no significant difference in apoptotic cell death in WT or Nan/+ FL cells (Fig. 7A and Supplementary Fig. S8A).

**Figure 7.**
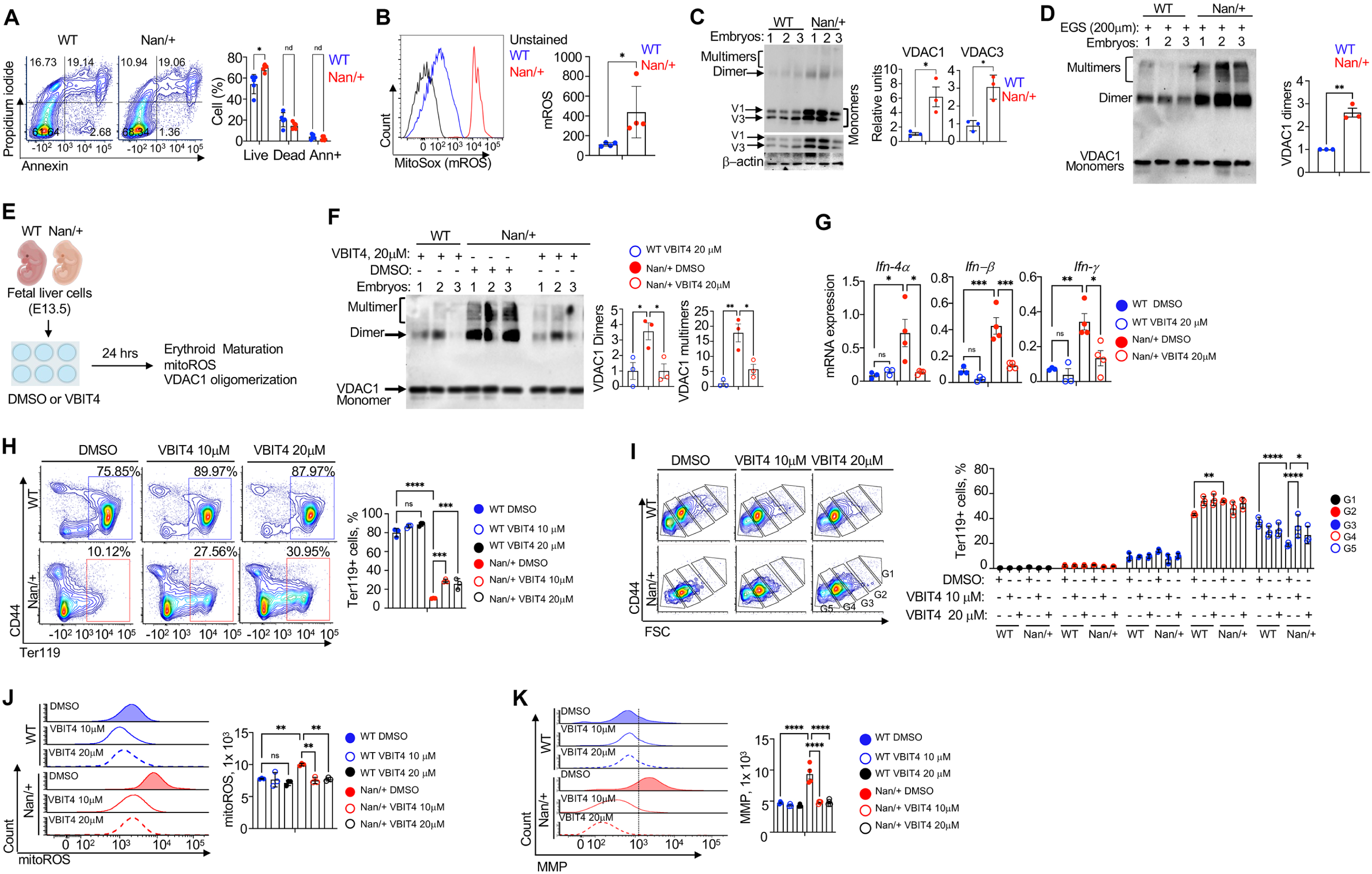
**VDAC oligomers facilitate release of mtDNA fragments and inhibit red cell maturation in Nan/+ FL cells** (A) Representative flow plot showing apoptotic cell death analyzed by staining with annexin V-FITC/propidium iodide (left) in WT or Nan/+ FL cells and their quantification (right). (B) Representative histogram of flow cytometric analysis (left) showing mitochondrial mitoSOX fluorescent ROS (mROS) measured by mitoSOX staining in FL cells obtained from WT or Nan/+. Quantitative analysis showing levels of mROS, as represented by Geo.mean (right). (C) Western blots of whole-cell extracts using an anti-VDAC1 antibody showing expression levels of VDAC1 (V1) and VDAC3 (V3) in freshly isolated FL cells from WT or Nan/+. VDAC oligomerization levels were shown without EGS-base cross-linking. VDAC monomers, dimers, and multimers are as indicated (left). VDAC1, VDAC3 relative units compared to β-actin. (D) VDAC1 oligomerization was visualized by immunoblotting using an anti-VDAC1 antibody after treatment with an EGS-based cross-linking reagent to stabilize the oligomers during electrophoresis in FL cells obtained from WT or Nan/+. The positions of VDAC1 monomers, dimers, and multimers are as indicated (left) and quantification of VDAC1 dimers (right). (E) Schematic diagram showing VDAC inhibitor (VBIT 4, 20μM) 24 hr treatment workflow in FL cells (E13.5) derived from WT or Nan/+. (F) FL cells treated with VBIT4 for 24 hrs were analyzed for VDAC1 oligomerization after treatment with an EGS-based cross-linker to stabilize the oligomers, and quantification of dimers and multimers (**right**). (G) Relative mRNA levels of Ifn-4α, -ß, or -ψ FL cells treated with VBIT4 after 24 hrs. (H) Representative flow plot of FL cells treated for 24hrs with DMSO or VBIT4 (10 and 20 μM) ex vivo were analyzed for erythroid expression by CD44/Ter119 and quantification of Ter119+ % (right). (I) Representative flow plot of FL cells treated for 24 hrs with DMSO or VBIT4 (10 and 20 μM) were analyzed for red cell maturation and gated for G1-G5 as indicated (**Right**). (J) mitoROS analyzed in BM Ter119 + cells treated for 24 hrs with DMSO or VBIT 4 (10 and 20 μM); histograms (left) and quantification (right) represented by Geo.mean. (K) MMP analyzed in BM Ter119 + cells treated with 24 hrs with DMSO or VBIT 4 (10 and 20 μM; histograms (left) and quantification (right) represented by Geo.mean. Each experiment is an average for *ex vivo* (n=3) and *in vivo* (n=5 to 6). In all panels, data are presented as mean ± S.E.M. (^∗^p < 0.05, ^∗∗^p < 0.01, and ^∗∗∗^p < 0.001, and ****p < 0.0001).

Alternatively, mitochondrial voltage-dependent anion channel 1 (VDAC1) is known to form oligomers under metabolic or mitochondrial oxidative stress conditions (mROS), and these oligomers can form large mitochondrial outer membrane pores ^60,64^. VDAC is the most abundant protein in the mitochondrial outer membrane and is a multifunctional channel that helps in regulating Ca2+ influx, metabolism, inflammasome activation, and apoptosis ^65,66^. mROS is an important trigger for increases in VDAC oligomerization ^60,67^. Increases in mitochondrial metabolism trigger mitochondrial stress, which can be analyzed by measuring the levels of mitochondrial ROS (mROS); such stress is enough to promote VDAC oligomerization. As a result, we measured mROS production using the MitoSOX fluorometric dye and find that Nan/+FL cells show significantly enhanced production of mROS (Fig. 7B and Supplementary Fig. S8B).

The expression levels of VDAC1, the most abundant of the three VDAC isoforms ^65^, and VDAC3 are elevated in autoimmune disease and, in some models, VDAC3 activity is thought to be triggered by released mtDNA ^60,68^. VDAC1 dimers, trimers, tetramers, and multimers can be readily assessed in cell extracts (Fig. 7C and Supplementary Fig. S8C). We find elevated quantities of VDAC1 and VDAC3 at both protein and mRNA levels in Nan/+ FL cells (Fig. 7C and Supplementary Fig. S8D). These data raise the possibility that the damaged mitochondrial cristate and increased VDAC levels may trigger the release of mtDNA to the cytosol. Evidence for VDAC1 oligomerization is enhanced when using chemical cross-linking agents followed by western blotting with anti-VDAC1 antibodies (Fig. 7D).

We propose mechanistically that oligomerized VDAC1 mediates mtDNA release to the cytosol, which in turn activates the mtDNA-cGAS-STING signaling pathway and induces a type-I IFN response in Nan/+ FL cells. These findings support the concept that abnormal processing and cytosolic release of mtDNA instigates intrinsic inflammatory signaling in Nan/+.

### Inhibition of VDAC1 oligomerization reduces inflammation in cultures improves anemia of Nan/+ mice in vivo

These findings suggest that oxidatively stressed mitochondria release mtDNA fragments via the channel formed by VDAC1 oligomers into the cytosol, thus inducing inflammation in Nan/+ FL cells. As a result, we tested the effect of the VDAC1 inhibitor, VBIT4, on the properties of the cells after 24h treatment of fresh E13.5 fetal liver cells in culture (Fig. 7E). VDAC1 oligomerization is decreased after treatment (Fig. 7F), and expression of inflammatory genes such as the interferons and cytokines/chemokines revert back to near WT levels (Fig. 7G, and Supplementary Fig. S8E). At the cell level, VBIT4 treatment increased red cell maturation as judged by Ter119+ recovery (Fig. 7H,I), and significantly decreased oxidative stress as measured by lower mtROS production (Fig. 7J) and MMP levels (Fig. 7K).

These results led us to test the effect of VBIT4 and another inhibitor, VBIT12, *in vivo* in adult mice by gavage treatments every two weeks, followed by harvest and analysis of bone marrow and peripheral blood (Fig. 8A). There is a high level of specificity, as only VBIT4 treatment significantly improves anemic hallmarks and ineffective erythropoiesis, results most readily apparent after the third treatment (6 weeks) as judged by elevated RBC counts, hematocrit and reduced reticulocytes (Fig. 8A). Consistent with our ex vivo results, BM Ter119+ cells are moderately but significantly elevated (Fig. 8B) and erythroid cell maturation is increased (Fig. 8C). Furthermore, mtROS levels (Fig. 8D) and MMP activity (Fig. 8E) are decreased in BM Ter119+ cells from VBIT4-treated Nan/+ mice compared to WT or DMSO-treated controls. We also find that IFN protein levels in serum are reduced in Nan/+ mice treated with VBIT4 (Fig. 8F). Finally, inhibiting VDAC oligomerization reduces splenomegaly in Nan/+ mice (Fig. 8G). Collectively these results demonstrate that treatment directed at mtDNA release/VDAC activity reduces inflammation *ex vivo* and *in vivo* in the adult Nan/+ mouse.

**Figure 8.**
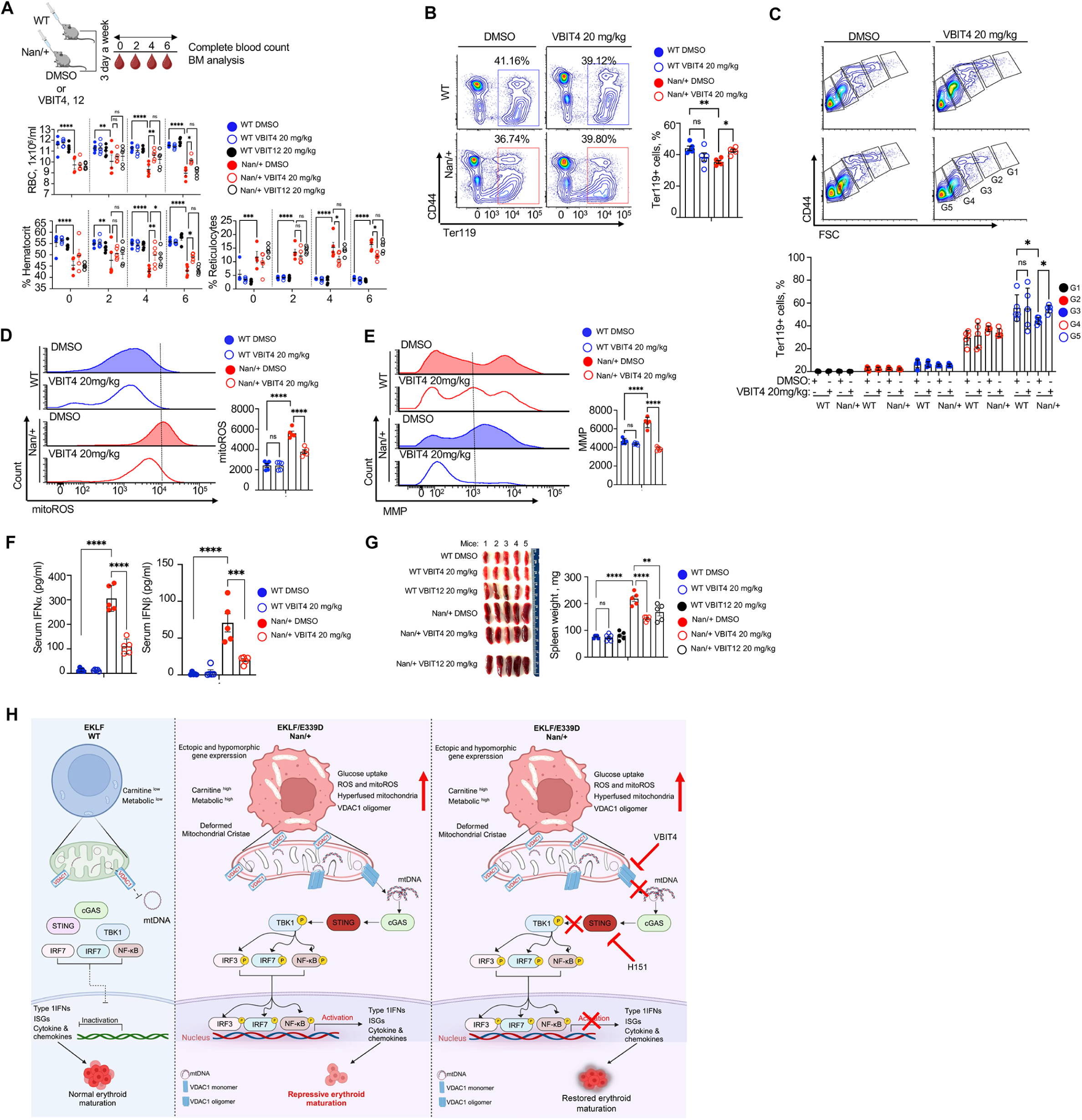
**VDAC1 inhibitor treatments significantly improves anemic hallmarks in adult Nan/+ mice.** (A) Schematic diagram showing VDAC inhibitor (DMSO or VBIT4 or VBIT 12, 5 mg/kg) treatment. Oral gavage was used to introduce mice with DMSO (control) or VBIT4 or 12, and blood samples analyzed at various weeks as indicated (left top), followed by total harvest of bone marrow (BM) at the end for blood parameter. Quantification of RBC numbers (top middle), hematocrit (left bottom) and reticulocyte counts (right bottom). (B) Representative flow plot of BM cells isolated from DMSO or VBIT 4-treated mice and were analyzed for Ter119+ (left) and quantification (right). (C) Representative flow plot of bone marrow cells isolated from DMSO or VBIT 4-treated mice and were analyzed for red cell maturation and gated for G1-G5 as indicated (bottom), and quantification (bottom). (D) mitoROS analyzed in BM Ter119 + cells treated with DMSO or VBIT 4; histograms (left) and quantification (right) represented by Geo.mean. (E) MMP analyzed in BM Ter119 + cells treated with DMSO or VBIT 4; histograms (left) and quantification (right) represented by Geo.mean. (F) Serum IFN-α and -ß levels measured using ELISA in mice treated with DMSO or VBIT4. (G) Spleens were isolated from DMSO or VBIT4- or VBIT12-treated mice and weighed. Size is shown (**left**) and weight is quantified (**right**). Each experiment is an average (n=5). In all panels, data are presented as mean ± S.E.M. (^∗^p < 0.05, ^∗∗^p < 0.01, and ^∗∗∗^p < 0.001, and ****p < 0.0001). (H) *Model*: Nan/+ fetal livers (E13.5) show ectopic and hypomorphic gene expression patterns due to the presence of the E339D variant. These cells are aberrantly enriched in L-carnitine and other metabolites. Their mitochondrial morphology is hyperfused and exhibit enhanced, but variable, degrees of disorganization, such as loss of cristae and detachment of inner membrane from outer membrane. Oligomerized VDAC1 mediates mtDNA release to the cytosol, which activates the mtDNA-cGAS-STING signaling pathway leading to increased transcripts for IFNs, interferon response genes (ISGs), and several inflammatory chemokine and cytokines. This mitochondrial dysregulation induces an inflammatory environment that inhibits proper erythroid maturation and causes ineffective erythropoiesis in Nan/+ embryos which persists into adulthood. Inhibition of the mtDNA-cGAS-STING module such as VDAC and STING with small molecules lowers the inflammatory response specifically in the Nan/+ erythroid cells and restores ineffective erythropoiesis seen in embryos fetal liver cells and in adults. (Model drawn via Biorender).

## DISCUSSION

Our studies demonstrate that the presence of mutant Nan-EKLF transcription factor, even when expressed as a heterozygote, has myriad metabolic consequences that logically flow from previously identified hypomorphic and neomorphic changes in downstream gene expression targets ^30–32^. Perusal of these strongly inferred that alteration of cell metabolism must follow and likely plays a contributing role in the neonatal anemia. We have shown just how pervasive such effects are, as they encompass direct effects on amino acid and nucleotide catabolism/anabolism, energy/ATP production, glucose utilization, and fatty acid synthesis/oxidation. The dysmorphia goes deeper, as it also includes altered mitochondrial structure/function. Mitochondrial stress induces mtDNA leakage into the cytosol in Nan/+ FL cells and activates the mtDNA-cGAS-STING pathway, leading to inflammation via a type I IFN response. In this context, we found mechanistically that oligomerized VDAC1 is responsible for mediating mtDNA release to the cytosol and inhibiting either the cGAS-STING pathway or VDAC1 oligomerization in the Nan/+ erythroid cell can restore normal red cell maturation. These observations (summarized in Fig. 8H) make it remarkable that the red cell endures such an onslaught of deconstructed pathways and that the animals survive until adulthood.

The metabolic changes in Nan/+ are shown to occur during early development (analysis of fetal liver erythroid cells) and to continue in the adult (analysis of CBC as well as BP). As a result, both intrinsic red cell changes as well as systemic effects follow from what is a single conservative amino acid substitution in one allele of EKLF.

Our previous studies had shown that expression of ectopic targets leads to systemic effects that followed from secreted factors that red cells normally do not produce ^32^. In the present analysis, the altered metabolome in the Nan/+ FL cells is consistent with accumulation of inflammatory molecules following initiation of the cGAS-STING pathway whose downstream effects negatively impact mitochondrial function (summarized in Fig. 8H). This likely contributes to the general inhibition of erythropoiesis and anemia in the Nan/+ mouse, leading to splenomegaly and increased circulatory levels of erythropoietin ^32^. At the same time, inflammation is known to stimulate stress erythropoiesis as part of a coordinated response to organismal insult ^3^. Typically, erythroid expansion of stress erythroid progenitors relies on glucose and favors anabolic processes to enable efficient cell growth and division. However, although we see enhanced levels of some amino acids in adult blood cells, this is not true across the board (e.g.,nucleotides) in Nan/+ FL cells. As a result, our observations do not appear to precisely fall into this scenario of inflammation/stress. Consistent with this, it has been noted that the stress erythropoiesis is significantly hampered in Nan/+ adults ^31^.

The most direct relevance of our observations to human erythroid biology is with CDA type IV (OMIM 613673), which follows from a heterozygous mutation at the same amino acid as in Nan/+, albeit a different substitution (E to K (CDA/IV) rather than E to D (Nan)) ^69^. These rare patients are highly anemic and transfusion dependent and present with morphologically aberrant red blood cells that exhibit an altered expression pattern mimicking that seen in Nan/+, including neomorphic gene expression ^70^. It is anticipated that these patients will likely also exhibit aberrant changes in their erythroid metabolome that will also have system effects. In support of this idea, a recent proteomic analysis of a human line (BEL-A) containing the CDA/IV mutation demonstrates that these cells exhibit increases in mitochondrial biogenesis, in components of the electron transport chain, and in the cristae organizing system ^71^. The proteomics data also shows a significant elevation in the STING protein level in CDA IV BEL-A cells attesting to the human disease relevance of our findings from the Nan/+ mouse.

Relatedly, inheritance of compound heterozygous EKLF mutants, though individually benign, can lead to problems; e.g., pyruvate kinase deficiency in humans ^72^. Given the role of this critical metabolite in proper red cell function and systemic health ^73,74^, our studies suggest that aberrancies in other EKLF-dependent metabolic pathways will be illuminated by analysis of red cells from these patients and others who inherit two imperfect copies of EKLF.

The increased levels of FAs in Nan/+ FL cells we have discovered here, including short chain fatty acids (SCFA) such as β-hydroxybutyrate (βHB), are likely to affect erythroid function in at least three ways. One is that changes in levels of acylated histones ^75,76^, which can be covalently modified by butyrate or ßHB ^76^, can have an effect on downstream gene expression ^77,78^. Two is that SCFAs are known inhibitors of HDAC proteins ^79–81^. EKLF is acetylated by P300/CBP ^82^, and its modification status plays a role in its optimal activity by enabling interactions with chromatin remodeling proteins ^83^. As a result, it is easy to envision that changes in expression of SCFAs can eventually lead to altered EKLF function in transcription/chromatin. Three, in humans, SCFAs are known stimulators of γ-globin expression ^84–86^, and thus have been touted as potential remedies for hemoglobinopathies such as ß-thalassemia and Sickle cell disease. As a result, these intracellular changes may be at least partially contributing to changes in hemoglobin switching capabilities observed in the Nan/+ mouse ^27^ and in human CDA/IV patients ^69^.

## ACKNOWLEDGEMENTS

This work was supported by NIH/NIDDK R01 DK046865 and by a Cooperative Center of Excellence in Hematology Pilot and Feasibility award (NIDDK). We thank the Flow Cytometry, Microscopy, Genomics, and Minerva HPC Cores at the Icahn School of Medicine at Mount Sinai for technical help. Electron microscopy tissue preparation and imaging were performed at the Microscopy and Advanced Bioimaging CoRE at the Icahn School of Medicine at Mount Sinai. We thank Varda Shoshan-Barmatz and Anna Kuzmine-Shteinfer (Ben Gurion University, Israel) for advice on the VDAC oligomerization protocol. We acknowledge Sanjana Pillay and Antanas Planutis for ongoing discussions.

## Authorship Contribution

T.A., and J.J.B. designed experiments, T.A., K.M., and L.X. performed experiments and T.A., K.M., analyzed data. T.A., K.M., and J.J.B. wrote the manuscript. All authors approved the manuscript.

## METHODS

### Data availability

The original LC-MS and GC-MS metabolite data, and results of the metabolomics analyses can be found in Data Source 1, 2 and Supplementary Tables S1-S8. Gene expression data and analyses are in Supplementary Tables S9-S15. Primer sequences are listed in Supplementary Table S16.

### Reagents and Resources

All reagents, kits, chemicals, and software are listed in Supplementary Table S17.

### Mice

Previously established *Nan* mouse strain (*Klf1^Nan^*/*Klf1^+^*) ^27–29,87^ were used for all experiments in this study. Because there is no significant difference in hematologic parameters among sexes ^27^, we did not perform sex genotyping on the embryos. Heterozygous adult Nan mice are termed Nan/+, and their strain is totally inborn (C3H/101 background). Embryos are produced by mating males with the Nan/+ allele to females with the +/+ allele. At embryonic day 13.5, mice were euthanized. All mouse breeding and animal experiments were approved by the Institutional Animal Care and Use committee under protocol IACUC-2019-0042 at Mount Sinai.

### Circulatory Blood Cell, Plasma, and Fetal Liver Cell Isolation

All animals were raised at 21 ± 2 °C with a 12-hour light/dark cycle. Blood from WT or Nan/+ adult mice, 200 μL of mandibular vein blood was drawn into a heparinized tube. Blood samples were centrifuged for 10 minutes at 5000×g and 4 °C. Supernatant blood plasma (BP) was isolated and immediately stored in 50 μL aliquots at -80 °C until analysis. 50 μL of circulatory blood cells (CBC) pellet were collected and washed with 200 μL NaCL 0.9%. The blood cell pellets were immediately kept in 50 μL aliquots at −80 °C until evaluation.

Embryonic fetal liver (FL) cells were isolated as described previously ^88^. Briefly, FL at E13.5 were dissected from WT or Nan/+ littermates and were homogenized into single cells by gentle pipetting and straining through a 70μM filter into ice-cold PBS containing 2% FBS.

### Liquid Chromatography-Mass Spectrometry (LC-MS) Metabolomics Sample Preparation

Isolated blood plasma (25 µL) from WT or Nan/+ as described above were thawed at 4°C for 45 minutes, then extracted with 75 µL of ice-cold acetonitrile (ACN,) + 0.1% ammonium hydroxide (NH_4_OH) containing internal standards and vortexed for 30 s. Cell pellets were extracted with 100 µL ice-cold 3:1 ACN:ddH_2_O + 0.1% NH_4_OH containing internal standards. Samples were then vortexed for 30 s after which 10 µL was aliquoted for acyl-carnitine analysis.

For CBC and FL cells from WT and Nan/+ isolated as described above were extracted with 450 µL of ice-cold ddH_2_O + 0.1% NH_4_OH containing internal standards, vortexed for 30 s, then 25 µL of reconstituted erythrocytes was taken forward for extraction. All samples were spiked with internal standards of 0.1 µg/mL of carnitine-D9) and 1 µg/mL tyrosine-D4. For all samples a process blank (PB) was created containing only extraction solvent, internal standards and ddH_2_O, and was carried through all extraction steps. After extraction and vertexing, all samples were chilled for 1 hour at -20°C, centrifuged for 10 minutes at 20,000 rcf at 4^O^C then transferred to new PTFE autosampler vials (Agilent) and injected in a randomized order. For all biological sources, a QC sample was prepared by pooling 10 µL from each sample.

### LC-MS Metabolomics Analysis

#### LC-MS General Metabolomics

A SCIEX 6500 QTRAP coupled to a SCIEX Nexera HPLC system (SCIEX, Framingham, MA USA) in both positive- and negative-modes was used for analysis. Separation was achieved using a Sequant Zic-pHILIC 2.1 x 100 mm column (Millipore Sigma, Burlington, MA, USA) with Phenomenex Krudkatcher (Phenomenex, Torrence, CA, USA). An initial concentration of 99% ACN with 5% ddH_2_O (buffer B) and 5% 25 mM ammonium carbonate (NH_4_)2CO_3_) in ddH_2_O (buffer B) was held for 1 min at a flow rate of 0.15 mL/minute. Buffer B was decreased to 15% over 14 minutes and further decreased to 5% over 4 minutes. Buffer B was returned to starting conditions over 0.1 minutes, and the system was allowed to re-equilibrate for 10 minutes between runs at a flow rate of 0.15 mL/minutes.

#### LC-MS Acyl-CoAs

A SCIEX 6500 QTRAP coupled to a SCIEX Nexera HPLC system in negative-mode was used for analysis. Separation was achieved using a Waters BEH amide 50 x 2.1 mm column (Waters, Milford, MA USA) with Phenomenex Krudkatcher Ultra. An initial concentration of 99% ACN with 5% ddH2O (buffer B) and 5% 25 mM (NH_4_)2CO_3_ in ddH2O (buffer B) was held for 0.5 minutes at a flow rate of 0.4 mL/minutes. Buffer B was decreased to 5% over 4.5 minutes. Buffer B was returned to starting conditions over 0.1 minutes, and the system was allowed to re-equilibrate for 5 minutes between runs.

### Analysis of LC-MS Mass Spectrometry Data

Data was collected using SCIEX analyst. Chromatogram integration was performed using SCIEX MultiQuant. This data was transferred to an Excel spread sheet (Microsoft, Redmond WA). Statistical analysis was performed using Microsoft Excel Data Analysis add-in. Metabolite identity was established using a combination of an in-house metabolite library developed using pure purchased standards and the METLIN library (https://metlin.scripps.edu/landing_page.php?pgcontent=mainPage). 3-indolepropionate, 4-aminobutyrate, 5-aminopentanoic acid, 5HIAA, biotin, choline, ethanolamine phosphate, homoserine, metanephrine, porphobilinogen, pyridoxamine, S-lactoylglutathione, sn-Glycero-3-phosphocholine, and theanine were verified by precursor and product ion only. All other metabolite identities were confirmed by MRM as well as retention time.

### Acyl Carnitine Sample Preparation

Acyl carnitines from plasma, circulatory blood and FL cells from WT or Nan/+ were extracted using a biphasic solvent system of cold methanol, methyl *tert*-butyl ether (MTBE), and PBS/water (See reference below) with some modifications. In a randomized sequence, to each sample was added 225 µL MeOH with internal standards and 750 µL MTBE. Samples were sonicated for 60 seconds and then incubated on ice with occasional vortexing for 1 hour. 188 µL of PBS was then added to induce phase separation. Vortexed for 10 seconds, rested at room temperature for 15 minutes, and then centrifuged at 15,000 x g for 10 minutes at 4 °C. The organic (upper) layer was collected, and the aqueous (lower) layer was re-extracted with 1 mL of 10:3:2.5 (*v/v/v*) MTBE/MeOH/dd-H2O, briefly vortexed, incubated at room temprature, and centrifuged at 15,000 x g for 10 minutes at 4 °C. Upper phases were combined and evaporated to dryness under speedvac. Lipid extracts were reconstituted in 250 µL of 1:1 (v/v) mobile phase A/B and transferred to an LC-MS vial for analysis. Concurrently, a process blank sample was prepared, and a pooled quality control (QC) sample was prepared by taking equal volumes (∼50 µL) from each sample after final resuspension ^89^.

### Acyl Carnitine LC-MS Analysis

Lipid extracted from plasma, circulatory blood and FL cells from WT or Nan/+ were separated on an Acquity UPLC CSH C18 column (2.1 x 50 mm; 1.7 µm) coupled to an Acquity UPLC CSH C18 VanGuard precolumn (5 × 2.1 mm; 1.7 µm) (Waters, Milford, MA) maintained at 60 °C connected to an Agilent HiP 1290 Sampler, Agilent 1290 Infinity pump, and Agilent 6490 triple quadrupole (QqQ) mass spectrometer. Lipids are detected using dynamic multiple reaction monitoring (dMRM) in positive ion mode. Source gas temperature is set to 200°C, with a gas (N_2_) flow of 14 L/minutes and a nebulizer pressure of 30 pounds per square inch (psi). Sheath gas temperature is 295°C, sheath gas (N_2_) flow of 12 L/minutes, capillary voltage is 3500 V, nozzle voltage 0 V, high pressure radio frequency (RF) 130 V and low pressure RF is 80 V. Injection volume is 1 µL and the samples are analyzed in a randomized order with the pooled QC sample injection eight times throughout the sample queue. Mobile phase A consists of H_2_O:ACN (98:2 *v/v*) in 10 mM ammonium formate (NH_4_HCO_2_) and 0.1% formic acid, and mobile phase B consists of IPA:ACN:H_2_O (90:9:1 *v/v/v*) in 10 mM NH_4_HCO_2_ and 0.1% formic acid. The chromatography gradient starts at 1% mobile phase B, holds for 1 minute then increases to 70% B over 8 minutes, increases to 99% B from 9-9.2 minutes where it’s held until 10.5 minutes then returned to starting conditions at 11 minutes. Post-time is 2.5 minutes and the flowrate is 0.5 mL/minute throughout. Collision energies and cell accelerator voltages were optimized using standards with dMRM transitions as [M+H]^+^→[*m/z* = 85.1] except for carnitine C0 which used [M+H]^+^→[*m/z* = 103 and 60.2].

### Gas Chromatography Mass Spectrometry (GC-MS)

#### Extraction of metabolite for GC-MS analysis

FL cells isolated from WT or Nan/+ as described above were mixed with cold 90% MeOH solution to give a final concentration of 80% MeOH to each FL cell pellet. Samples were then incubated at -20 ¼C for 1 hour. After incubation the samples were centrifuged at 20,000 x g for 10 minutes at 4 ¼C and supernatant was then transferred from each sample tube into a newly labeled micro centrifuge tube. Pooled quality control samples were made by removing a fraction of collected supernatant from each sample and process blanks were made using only extraction solvent and no cell culture. The samples were then dried *en vacuo*.

#### GC-MS analysis of samples

All GC-MS analysis was performed with an Agilent 5977b GC-MS MSD-HES fit with an Agilent 7693A automatic liquid sampler. Dried samples were suspended in 40 µL of a 40 mg/mL O-methoxylamine hydrochloride (MOX) in dry pyridine and incubated for 1 hour at 37 °C in a sand bath. Then 25 µL of this solution was added to auto sampler vials. 60 µL of N-methyl-N-trimethylsilyltrifluoracetamide (MSTFA with 1% TMCS, (Trimethylchlorosilane) was added automatically via the auto sampler and incubated for 30 minutes at 37 °C. After incubation, samples were vortexed and 1 µL of the prepared sample was injected into the gas chromatograph inlet in the split mode with the inlet temperature held at 250 °C. A 5:1 split ratio was used for analysis for the majority of metabolites. Any metabolites that saturated the instrument at the 5:1 split were analyzed at a 50:1 split ratio. The gas chromatograph had an initial temperature of 60 °C for 1 minute followed by a 10 °C/min ramp to 325 °C and a hold time of 10 minutes. A 30-meter Agilent Zorbax DB-5MS with 10 m duraguard capillary column was employed for chromatographic separation. Helium) was used as the carrier gas at a rate of 1 mL/minute.

### Analysis of GC-MS Data

Data was collected using MassHunter software (Agilent). Metabolites were identified and their peak area was recorded using MassHunter Quant (Agilent). This data was transferred to an Excel spread sheet (Microsoft, Redmond WA). Metabolite identity was established using a combination of an in-house metabolite library developed using pure purchased standards, the NIST library and the Fiehn library.

### Statistical Analysis of LC-MS and GC-MS data

The raw LC-MS data was processed using locally weighted scatterplot smoothing, and then by normalization to the appropriate internal standard. Raw GC-MS data was normalized to internal standards and combined with the LC-MS data, when both LC-MS and GC-MS data sets are present. The data was checked for quality by assessing the percent coefficient of variation (%CV) of each metabolite in the pooled quality control samples. Metabolites with a %CV of > 30% were removed from the analysis. When overlap occurred between metabolites in LC-MS and GC-MS, the analysis with the lowest %CV was retained. Statistical analysis of the mass spectrometry data was performed using metaboanalyst.ca ^90^. The data was normalized by log-transformation and auto scaling. This was used at the input to generate PCA plots, heat maps, ANOVAs, t-tests, and volcano plots for the metabolomics data.

### Construction of Metabolic Pathway and Metabolic Set Enrichment Analysis (MSEA)

Metabolic pathway analysis and metabolic set enrichment analysis (MSEA) were performed using the MetaboAnalyst 5.0 platform and to identify the most significant metabolite that is altered in between WT and Nan/+ FL cells for identification of the top altered pathways and MSEA. We utilized KEGG IDs as input in MetaboAnalyst 5.0 pathway analysis tool by using flowing parameters for analysis. The Mus musculus library (version March2023) was chosen as the pathway library. We used hypergeometric test for metabolic pathway analysis and for MSEA analysis.

### Glucose uptake assay

For measurement of glucose uptake, freshly isolated FL cells (1x10^6^) from WT or Nan/+ were cultured immediately in 500μL of glucose, glutamine, pyruvate free medium containing 100 μM of 2-(n-(7-nitrobenz-2-oxa-1,3-diazol-4-ylamino)-2-deoxyglucose (2-NBD-Glucose or 2NBDG) for 2 hour after which cells were then washed three times in PBS, further re-suspended in PBS containing 1 μg/ml DAPI, and analyzed by flow cytometry LSR Fortessa Cell Analyzer (BD Biosciences) for 2NBDG fluorescence in the FITC channel. Quantification of 2NBDG uptake was measured by the geometric mean fluorescence intensity (MFI) as well as % of 2NBDG+ cells.

### ATP assay

Enzymatic measurement of ATP was carried out following the guidelines provided by the colorimetric ATP Assay Kit (Abcam). FL cells (5 x 106) obtained from WT, Nan/+, or adult liver mice were subjected to a PBS wash and subsequently lysed in 150 µL of ATP Assay buffer. Subsequently, the cells underwent centrifugation at 13,000 x g for 5 minutes at 4 °C to remove insoluble material. The resulting supernatant was then carefully transferred into a 96-well plate, with each well containing a volume of 50 µL in 3 technical replies and plate was incubated at room temperature for 30 minutes and protected from light. The absorbance was measured at wavelength λ = 570 nm by the Varioskan LUX multimode microplate reader (Thermo -Fisher Scientific). ATP concentration was standardized with the use of ATP solution of known concentration provide in the kit. The data were presented in terms of ATP concentration, measured in nanomoles per milligram of protein, along with the standard error of the mean (S.E.M.)

### mtDNA quantification

Total genomic DNA was isolated from FL cells (E13.5) were isolated from WT or Nan/+ using QIAamp DNA Micro kit according to kit instruction. DNA concentration was quantified by Thermo Scientific™ NanoDrop 2000 and 1 ng of DNA extract was loaded in the final reaction. qRT-PCR was performed using PowerUp SYBR® Green Master Mix and 7900HT ABI machine (see Primer sequences Supplemental Table S16). For mtDNA quantification, three sets of primers targeting the mitochondrially encoded cytochrome B (*MT-Cy B)* and mitochondrially encoded NADH dehydrogenase 4 (*MT-ND4)* genes and for nDNA quantificationprimers targeting the nuclear β-actin were selected for quantification of absolute copy number. Each DNA was generated from three independent biological replicates.

### RNA isolation and Real-time quantitative RT-PCR

Total RNA was isolated from WT or Nan/+ FL cells (E13.5) and using RNeasy Mini Kit (Qiagen) according to the manufacturer’s instructions. After DNase treatment, RNA was reverse transcribed using a SuperScript VILO cDNA synthesis kit according to the manufacturer’s instructions. RT-PCR was performed using RT² SYBR Green Fluor qPCR Mastermix in triplicates, using the indicated primers and 7900HT ABI machine. The relative level of each transcript was analyzed using to β-actin RNA levels as the housekeeping gene. Primer Sequences used in the study are in Supplemental Table S16.

### Measurement of plasma levels L-carnitine

L-carnitine were analyzed by using colorimetric/fluorometric assay kit according to manufacturer’s instruction. Briefly for the fluorescent assay 4 and 8 µl of deproteinized plasma isolated from WT or Nan/+ were directly diluted in the assay buffer to bring sample wells to 50 µl/well in a 96-well plate in technical triplicates. Deproteinization of plasma sample were performed using centrifugation through a 10kDa MW cut-off filter. Next, 50 µl of the reaction mix prepared according to the kit protocol were added to each well containing the carnitine standard, test and background control samples. Reaction mixtures were mixed thoroughly by pipetting and plate were incubated at room temperature for 30 min and protected from light. The absorbance was measured at Ex/Em 535/587 nm in a Varioskan LUX multimode microplate reader (Thermo Fisher Scientific). L-carnitine concentration was standardized with the use of L-carnitine solution of known concentration provide in the kit. The data were presented in terms of L-carnitine concentration, measured in nmol/µl, along with the standard error of the mean (S.E.M.) The normal range for serum L-carnitine is ∼ 10-70 µM.

### Measurement of plasma levels of β-hydroxybutyrate (BHB)

BHB levels in plasma were measured using a PicoProbe™ β-Hydroxybutyrate fluorometric assay kit following the manufacturers’ instructions. Briefly 4 and 8 µl of deproteinized plasma isolated from WT or Nan/+ were directly diluted in the β-Hydroxybutyrate assay buffer to bring sample wells to 50 µl/well in a 96-well plate in technical triplicates. Deproteinization of plasma sample were performed using centrifugation through a 10kDa MW cut-off filter. Next, 50 µl of the reaction mix prepared according to the kit protocol were added to each well containing the known standard, test and background control samples. Reaction mixtures were mixed thoroughly by pipetting and plate were incubated at room temperature for 30 min and protected from light. The absorbance was measured at Ex/Em 535/587 nm in a Varioskan LUX multimode microplate reader (Thermo Fisher Scientific). The data were presented in terms of L-carnitine concentration, measured in nmol/mL, along with the standard error of the mean (S.E.M.)

### Cytosolic reactive oxygen species (ROS), Mitochondrial ROS and Membrane potential measurements

Cytosolic reactive oxygen species (ROS) were analyzed in freshly isolated FL cells from WT or Nan/+. The cells were then processed with DCFDA / H2DCFDA - Cellular ROS assay kit according to the manufacturer’s instructions. Briefly, FL cells isolated from WT or Nan/+ were incubated in the working buffer for 30 minutes at 37 °C with 5% CO2 and then washed with HBSS twice. The cell pellet was resuspended in 500uL PBS and ROS level in cells was determined by flow cytometry LSR Fortessa Cell Analyzer (BD Biosciences). Data were analyzed with FlowJo 10.

According to the manufacturer’s instructions, mitochondrial ROS (mtROS) production was analyzed by using MitoSOX green a mitochondrial superoxide indicator. In brief, FL cells isolated from WT or Nan/+ were washed with warm PBS, treated with 2mM MitoSOX green at 37 °C for 45 minutes, and washed with warm PBS, resuspended in PBS (500 ml). Then, fluorescence (Excitation/Emission.

∼488/510 nm) was and data acquired by flow cytometry LSR Fortessa Cell Analyzer (BD Biosciences). Data were analyzed with FlowJo 10.

To measure mitochondrial membrane potential (MMP), Tetramethylrhodamine ethyl ester perchlorate (TMRE, 100nM) was used in accordance with the manufacturer’s instructions which specifically accumulates within the mitochondrial matrix of live cells. In brief, FL cells isolated from WT or Nan/+ were stained with the TMRE at 37 °C for 15 minutes. Flow cytometry acquisition was performed with a LSR Fortessa Cell Analyzer (BD Biosciences). All flow cytometry analyses and quantifications were performed using FlowJo 10 or FCS express software.

### Flow Cytometry

For the analysis of erythroblast populations, FL (E13.5) or bone marrow (BM) cells were obtained from WT of Nan/+ in IMDM + 2% FBS. Cells were filtered, washed with 2% FBS/PBS, and stained with TER119- pacific blue, CD44-FITIC, (1:100 dilution) in 2% FBS/PBS for 15 min at room temperature. After staining with primary antibodies cells were washed and finally resuspended in IMDM + 2% FBS containing DAPI (viability 1 µg/mL). For flow cytometry analysis-stained cells were acquired on the LSR LSR Fortessa and quantifications were performed using FCS express software.

### STING and VDAC inhibitors ex vivo treatment

In brief, FL cells from WT of Nan/+ (E13.5). The cells were washed with phosphate-buffered saline (PBS), resuspended in erythroid expansion culture media ^91^ and cultured in 12 well plates. Subsequently, the cells were treated with the DMSO (control) or with VBIT-4 (10 or 20 μM) for 24 hrs. After 24 hrs cells were washed and analyzed for erythroblast populations, VDAC1 oligomerization, mitochondrial ROS and mitochondrial activity (MMP) as described in the methods.

For sting inhibitor treatment FL cells from WT of Nan/+ (E13.5). The cells were washed with phosphate-buffered saline (PBS), resuspended in erythroid expansion culture media, and cultured in 12 well plates. Subsequently, the cells were treated with the DMSO (control) or with H151 or C176 (50 or 100 μM) for 24 to 72 hrs. At each time point cells were washed and analyzed for erythroblast populations and mitochondrial activity (MMP) as described in the methods.

### H-151 preparation and in vivo treatment

In brief, H-151 was initially diluted in DMSO to achieve a concentration of 20 mg/ml, and then further diluted in sterile phosphate-buffered saline (PBS) to reach a final DMSO concentration of 5% before being injected into mice. Aliquots of 2 ml were stored at -20 °C. Prior to injection, the mixture was allowed to warm to room temperature. All experiments used age-matched wild-type (WT) or Nan/+ mice, (aged between 10 and 12 weeks). Each mouse received H-151 at a dosage of 5 mg/kg every 2 days for a duration of 5 weeks via intraperitoneal injection. For complete blood composition (CBC), whole blood was drawn every two weeks, finally after 5 weeks of H151 treatment mice were killed and bone marrow was collected for further analysis.

### VBIT 4 and VBIT 12 preparation and in vivo treatment

VBIT-4 and VBIT12 was obtained from Selleckchem. VBIT-4 or VBIT12 was dissolved as previously described ^60,92^ and administered to WT or Nan/+ mice by oral gavage at a dose of 20 mg/kg body weight. Mice were oral gavage every second day for 6 weeks. Similar doses of VBIT-4 had been previously used in models of Alzheimer’s disease ^92^ and systemic lupus erythematosus ^60^. For complete blood composition (CBC), whole blood was drawn every two weeks, finally after 6 weeks of VBIT4 or VBIT12 treatment mice were killed, and bone marrow was collected for further analysis.

### Blood collection and Plasma IFNα or IFN β analysis

Blood was withdrawn from the retroorbital vein plexus from the mice treated with either H151 or VBIT4 (WT or Nan/+) and collected in with EDTA. Blood plasmas were collected by centrifuging at 850 g for 10 minutes at room temperature, and precipitated red and white blood cells were discarded.

Quantification of IFNα or IFN β content in the blood was performed on plasma using the Mouse IFNα or IFN β All Subtypes ELISA KIT High Sensitivity (pbl Assay Science) according to manufacturer’s instruction.

### CD71+ Ter119+ cell enrichment from mouse bone marrow

A fraction of the DMSO- and H151-treated mouse bone marrow cells (10^6^ cells) from WT and Nan/+ mice were used for erythroid cell isolation using the EZ-Sep Mouse PE positive selection kit. Cells were washed in 1X PBS and resuspended in 1X PBS + 2% FBS + 1mM EDTA followed by EZ-sep magnetic selection using the manufacturer’s protocol. The antibodies Ter119-PE and CD71-PE antibodies were each added at a 1:100 dilution to select for total PE+ cells at both early and mature stages of erythropoiesis. Two rounds of PE+ selection were performed and after selection, cells were washed and stored at -80°C for cell lysis and Western blot.

### Bone marrow Single-cell RNA-Seq

DMSO-treated and H151 treated WT and Nan/+ mice (n=3 each) were sacrificed, and total bone marrow was isolated from the hind limbs into 1X PBS + 2% FBS. Cells were filtered using a cell strainer and homogenized. 100,000 cells from each biological replicate were pooled for single cell sequencing. Libraries were generated using Chromium Single Cell Reagent Kit V3 (10X Genomics) to generate cDNA and barcoded indexes and paired-end sequencing was performed using a Novaseq instrument. Single-cell sequencing data was analyzed using the R package Seurat version 5 ^93^ with built-in functions for filtering and scaling, U-MAP clustering, cell-cycle scoring, expression plotting, differential expression analysis, and cell cycle scoring. The entire annotated R code along with the Seurat object used in our analysis will be deposited in Zenodo, and the raw and analyzed data will be deposited in the Gene Expression Omnibus (GEO) and will be publicly available as of the date of publication.

### Apoptosis assay

Annexin V-FITC/PI Apoptosis detection kit were used for analyzing cell apoptosis. In brief, FL cells (1x10^6^) isolated from WT or Nan/+ were washed with PBS, incubated with 500μl 1×binding buffer and stained with 5μl of Annexin V-FITC plus 5μl of PI. Cells were incubated for 20 minutes at room temperature in the dark. Finally, the fluorescence was immediately analyzed with a flow cytometer LSR Fortessa Cell Analyzer (BD Biosciences) within 1 hour.

### Electron microscopy

Electron microscopy experiments were performed as described ^94^. Briefly, freshly isolated FL cells from WT or Nan/+ were fixed with 2% paraformaldehyde/ 2.5% glutaraldehyde in 0.1M sodium cacodylate solution at 4°C for at least one week. Cells were rinsed 3 times with 0.1M sodium cacodylate buffer and fixed with 2% osmium tetroxide/1.5% potassium ferricyanide in 0.1M sodium cacodylate for 1 hour at room temperature, and *en bloc* stained with 2% uranyl acetate in dH20. Next, Cells were dehydrated in an ethanol series (25% up to 100%), infiltrated through an ascending ethanol/resin series and placed in resin overnight. Then samples were placed in a 60°C vacuum oven for 72 hours to polymerize. The blocks were ultrathin-sectioned (0.5 and 1 µm) were obtained using an ultramicrotome (Leica EM UC7; Leica Microsystems) and counterstained with 1% toluidine blue. Ultra-thin sections (80nms) were collected on copper 300 mesh grids using a coat-quick adhesive pen and all sections were counter-stained with 1% uranyl acetate and lead citrate. Images were taken on an HT7500 transmission electron microscope (Hitachi High-Technologies, Tokyo, Japan) using an AMT NanoSprint12 12megapixel CMOS TEM Camera (Advanced Microscopy Techniques, Danvers, MA). Final image brightness, contrast, and size were adjusted using Adobe Photoshop CS4 software version CS4 11.0.1 (Adobe, Inc., San Jose, CA, USA). Images of at least 30 mitochondria from at least 10 different cells were randomly collected at magnification of 1500× for each group. Deformed mitochondrial cristae morphology were quantified manually (visually).

### Chemical cross-linking

Chemical cross-linking was performed as described previously ^64^. In summary, FL cells from WT or Nan/+ were washed with PBS and then incubated with the cross-linking reagent EGS (200 μM) for 15 minutes at 30°C. 50 ug protein samples were subjected to SDS-PAGE and immunoblotting using anti-VDAC1 antibodies, as described below. Quantitative analysis of immuno-reactive VDAC1 dimer, trimer and multimer bands intensity was performed with Image J software.

### Extraction of nuclear/cytosolic fraction

The nuclear extraction was performed using a Nuclear/Cytosol fractionation kit in accordance with the guidelines provided by the manufacturer. In summary, the freshly isolated FL cells derived from WT or Nan/+ were washed 2 times in ice cold PBS and subsequently centrifuged at 600 g for 3 minutes. The pellet of cells was suspended in 200 μl of cytoplasmic extraction reagent A (CEB-A) by vigorous vertexing and incubated on ice for 10 minutes. Subsequently, 11 μl of a second cytosol extraction buffer-B (CEB-B) was added. The mixture was then vortexed for 5 seconds, followed by another incubation on ice for 1 minutes. Finally, the suspension was centrifuged at a speed of 16,000 x g for 5 minutes. Promptly transfer the supernatant fraction containing cytoplasmic extract into a pre-cooled tube. Next, the insoluble pellet fraction, which consisted of crude nuclei, was resuspended in 100 μl of nuclear extraction buffer by vertexing for 15 seconds. This resuspended mixture was then incubated on ice for a period of 40 minutes, followed by centrifugation at 16000xg for 10 minutes. The supernatant, which contained the nuclear extract, was stored at a temperature of -80°C for future utilization and was employed in the subsequent experiments.

### Immunoblot analysis

Freshly isolated FL cells from WT or Nan/+ were lysed using 1 x cell lysis buffer supplemented with protease inhibitor cocktail inhibitors. Next, supernatant was mixed with a 5x protein loading buffer. The protein lysates were boiled at 60 °C for 5 minutes, and the protein concentration were determined using a Barford assay. Aliquots of 15–30 μg protein per sample were fractionated on 10% SDS-PAGE gels, transferred onto nitrocellulose membranes, and with 5% non-fat dry milk and 0.1% Tween-20 in Tris-buffered saline. Membranes were incubated with primary antibodies (1:1000) overnight at 4 °C, extensively washed, and incubated with HRP-conjugated anti-mouse or anti-rabbit whole IgG secondary antibodies (1:5000) for 3 hours at room temperature. Western Bright ECL HRP substrate was then applied, and protein bands were detected using a ChemiDoc MP system (BioRad Laboratories, USA). Protein band intensity quantification analysis was performed with Image J software. The antibodies that were used for western blots are listed in the star methods.

### Immunofluorescence Staining, Imaging and Analysis Laser scanning confocal microscopy

Immunofluorescence Staining was performed as described previously ^47^. In brief, FL cells (1x10^4^) isolated from WT or Nan/+ were fixed with 4% paraformaldehyde for 15 minutes. Cells were washed three times with PBS and permeabilized in PBS + 0.25% Triton X100 for 20 minutes and blocked for 2 hours in 3% BSA at room temperature. Next, fixed and permeabilized cells were then incubated with primary antibodies (1:100) overnight at 4°C. After incubation cells were washed and stained with fluorescence-conjugated secondary antibodies (1:1000) for 2 hours at room temperature. Finally, slides were sealed with mounting medium with DAPI. Images were captured using a Zeiss LSM880 Airyscan confocal microscope using a 100 X objective (N.A. 1.46).

### Super resolution confocal microscopy

All images were collected using a Zeiss LSM 880 confocal microscope with Airyscan Super Resolution Imaging module and a 100X/1.46 Alpha Plan Apochromat objective lens (Zeiss MicroImaging, Jena, Germany) with "optimal" (Nyquist) XY scaling as described previously ^47^. Z stacks over the whole of the cell were taken at a resolution of 0.018 millimeters (at least 25 optical sections), a pixel dwell period of more than 50 microseconds, and field dimensions of 300 millimeters on a side. Total of at least 20 WT and 20 Nan/+ FL cells were analyzed. This was then followed by the processing of images using Airyscan (set to auto but seldom over 6.2), and analysis using ZEN image capture and processing software (ZEN blue/black). Additionally, the highest-intensity projections that are shown in the figures were acquired with the help of the ZEN Blue program.

### Image analysis

Unless otherwise noted, all images were analyzed using either the FIJI or NIH ImageJ software ^95^. The brightness and contrast levels were determined during the image capturing process and were not changed during processing.

### Fluorescence intensity

The protein of interest was isolated, and its labeling was quantified on a per cell basis using the raw integrated intensity metric obtained from the measure command. For measuring nuclear intensity, the nuclei were delineated using DAPI thresholds and then applied to the channel showing the protein of interest. This allowed for the measurement of fluorescence intensity exclusively within the nucleus.

### Mitochondrial morphology

Freshly isolated FL cells from WT or Nan/+ were analyzed for mitochondrial morphology as described previously ^47^. In brief, Imaris software (for 3D images) was used to do an individual analysis on each cell in order to identify the degree to which mitochondrial fragmentation had occurred. Using the Imaris program, the surface of the mitochondria was first produced for the 3D images. After that, the total number of pieces as well as the total area of the surface were measured. For Mitochondrial morphology, more than 30 cells were studied for WT and Nan/+.

### Colocalization

The cells were manually chosen, and the channels that contained the two proteins of interest were isolated and examined using the Colocalization plugin in Fiji software. A total of at least 30 cells from WT or Nan/+ FL cells were analyzed. The Coloc function automatically determines the threshold and provides a calculated value for Mander’s correlation coefficients. The degree of colocalization between two proteins was assessed by calculating the average across all cells examined within each group. The calculation of colocalization percentage was performed using the JACoP plugin, which is a tool available in the NIH Fiji software.

### Quantification and statistical analysis

Unless indicated elsewhere, all statistical analysis were completed in GraphPad Prism 9. Data is presented as mean ± S.E.M. One-way ANOVA was used for comparison across more than two groups. The specific statistical test used for each specific experiment is indicated in each figure legend. All experiments were performed at least with three biological replicates unless specified. p < 0.05 was considered significant in all experiments p< 0.05, ^∗∗^p < 0.01, ^∗∗∗^p < 0.001, ^∗∗∗∗^p < 0.0001 and n.d (non-significant).

