## Supplemental Figures S1-S8 for "STING and VDAC inhibitors attenuate inflammation and ineffective erythropoiesis caused by an altered metabolome in the *Nan* (EKLF/E339D) mouse model of neonatal anemia"

### Figure S1

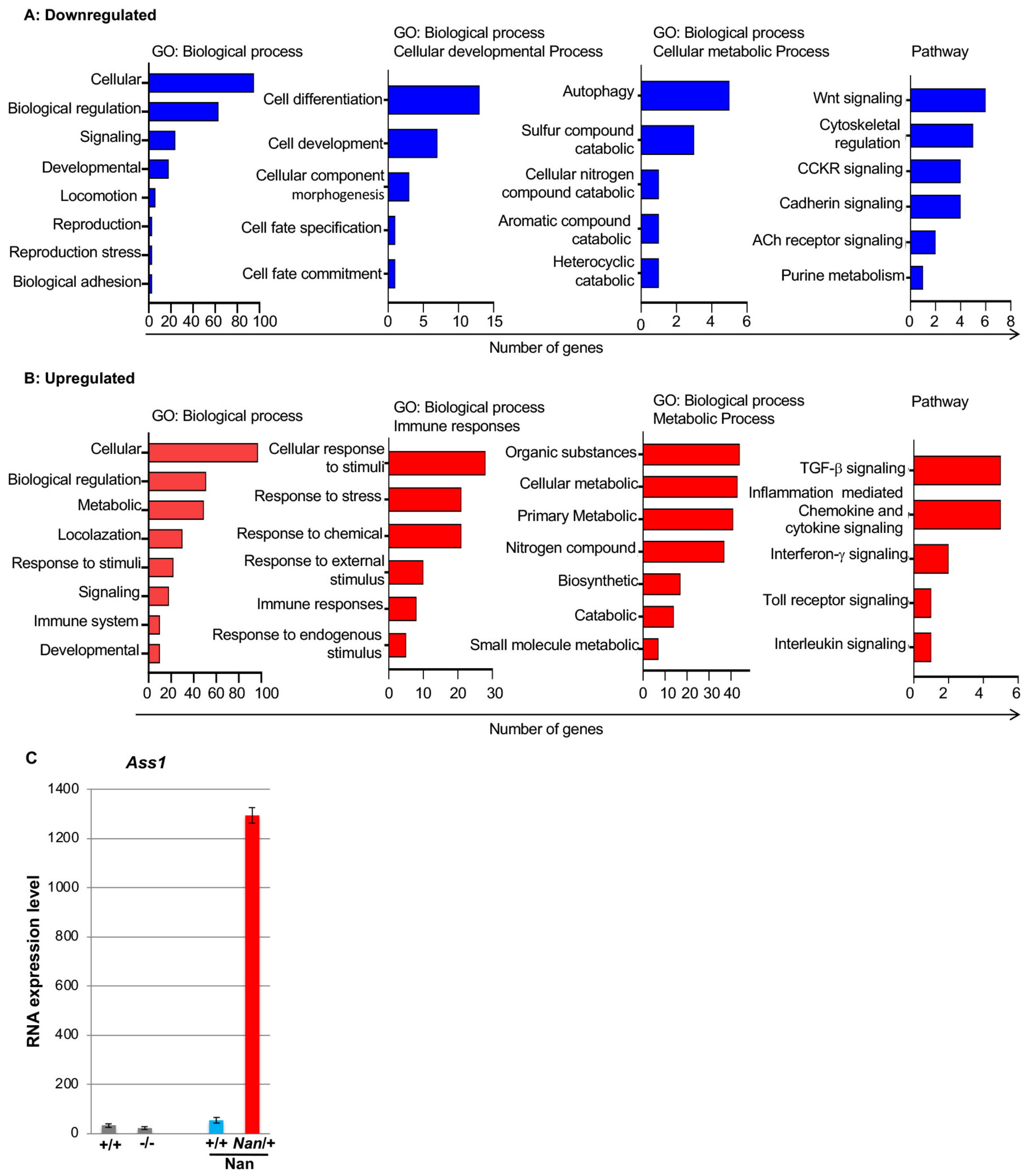


**Figure S1. Ectopic/neomorphic gene expression in the Nan/+ mutant fetal liver cell**
Gene ontology and pathway analysis of differential gene expression between WT and Nan/+ FL cells from previously performed bulk RNA sequencing (ref 33). PANTHER enrichment pathway analysis of downregulated **(A)** and upregulated **(B)** genes in WT vs. Nan/+ FL cells are as indicated. The top items of biological processes and pathways using the PANTHER database are shown. The identified pathways are represented on the y axis; the x-axis represents the number of significant genes found in each pathway. Data are presented from n=3 biological replicates per group. **(C)** RNA-seq expression data of E13.5 fetal liver cells from WT (+/+) and EKLF-null (-/-) littermates (left), or from WT (+/+) and Nan/+ littermates ("Nan") (right), were analyzed for expression of argininoosuccinate synthetase (Ass1) (samples are from refs 9,33). Each point is an average of biological triplicate samples.

### Figure S2

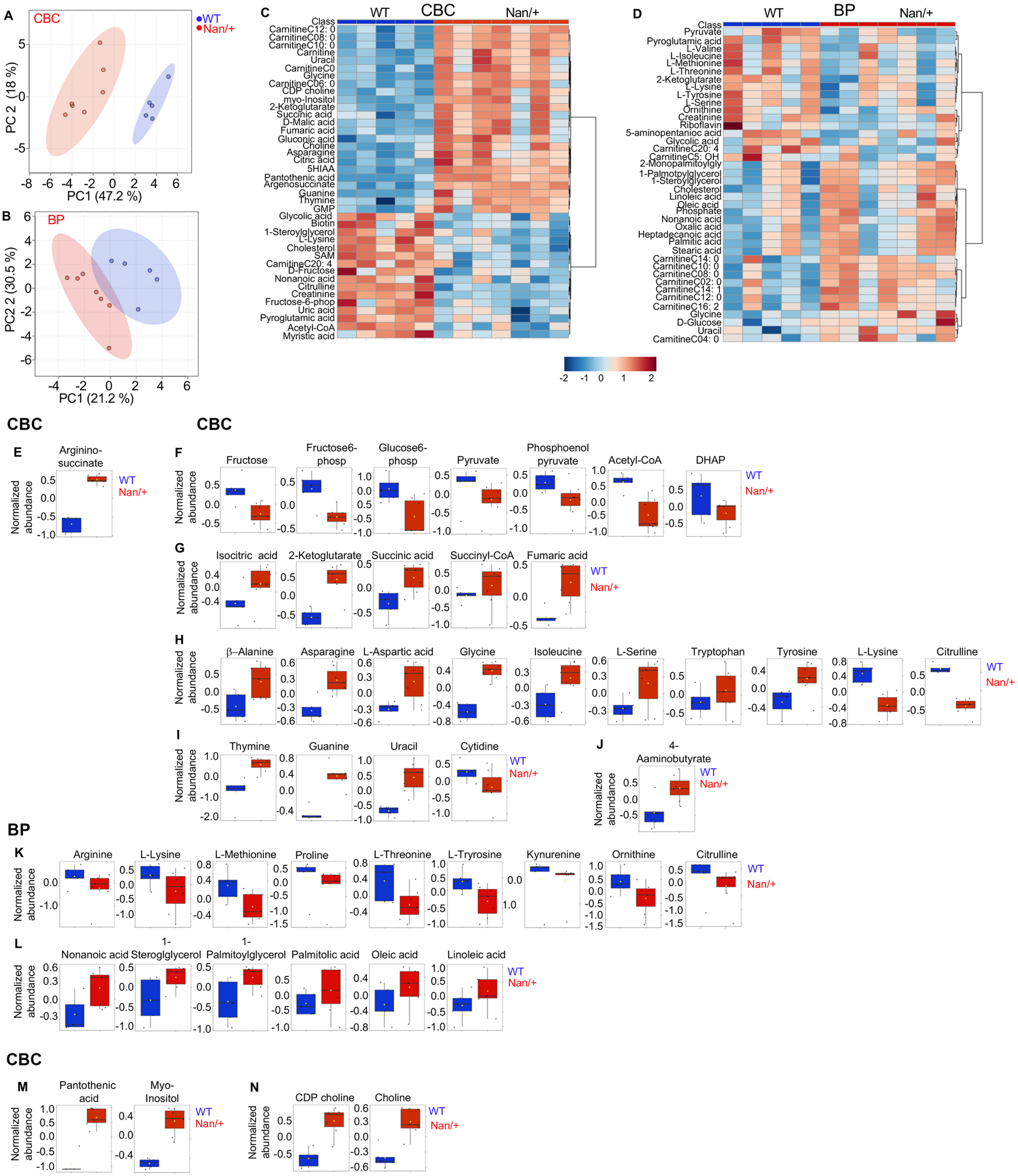

A
CBC
WT
Nan/+
PC 2 (18 %)
PC1 (47.2 %)
B
BP
PC2 2 (30.5 %)
PC1 (21.2 %)
C
WT
CBC
Nan/+
Class
D
WT
BP
Nan/+
Class
CBC
CBC
E
Arginino-succinate
F
Fructose
Fructose6-phosp
Glucose6-phosp
Pyruvate
Phosphoenol pyruvate
Acetyl-CoA
DHAP
G
Isocitric acid
2-Ketoglutarate
Succinic acid
Succinyl-CoA
Fumaric acid
H
β–Alanine
Asparagine
L-Aspartic acid
Glycine
Isoleucine
L-Serine
Tryptophan
Tyrosine
L-Lysine
Citrulline
I
Thymine
Guanine
Uracil
Cytidine
J
4-Aaminobutyrate
BP
K
Arginine
L-Lysine
L-Methionine
Proline
L-Threonine
L-Tryrosine
Kynurenine
Ornithine
Citrulline
L
Nonanoic acid
1-Steroglglycerol
1-Palmitoylglycerol
Palmitolic acid
Oleic acid
Linoleic acid
CBC
M
Pantothenic acid
Myo-Inositol
N
CDP choline
Choline
Normalized abundance

**Figure S2. Global profiling of late developmental circulatory blood cells and plasma shows significant differences in a wide range of compounds in Nan/+**

**(A-B)** Principal component analysis (PCA) plot shows analysis from adult WT or Nan/+ circulatory blood cells (CBC) (**A**) and blood plasma (BP) (**B**) (WT in blue, Nan/+ in red). As is evident in both groups, WT data points are well separated from Nan/+.

**(C-D)** The heat map (distance measure using euclidean, and clustering algorithm using ward. Heat map depicts the top 40 metabolites obtained in analysis that are unique to WT or Nan/+ in CBC **(C)** and BP **(D).**

**(E)** Box plot showing relative abundances of arginosuccinate in CBC obtained from WT or Nan/+ adult mice (p=2.61E-07).

**(F-I)** Box plot showing relative abundances of glycolysis **(F),** mitochondrial TCA cycle **(G),** amino acid **(H),** and nucleotide **(I)** related metabolites from WT or Nan/+ CBC from adult mice (p<0.05; ranges= 0.049882 to 1.70E-07).

**(J)** Box plot showing relative abundances of 4-aminobutyrate in CBC obtained from WT or Nan/+ adult mice (p=2.61E-07).

**(K-L)** Box plot showing relative abundances of amino acid **(K)** and several fatty acid (**L)** related metabolites in the BP of adult WT or Nan/+ mice (p<0.05; ranges= 0.042103 to 0.00043188).

**(M-N)** Box plot showing relative abundances of pantothenic acid (p=7.79E-06) and myo-insitol (p=6.06E-05) **(M)** and CDP choline (p= 0.00033376) **(N)** related metabolites in CBC obtained from WT or Nan/+ adult mice.

For **(E-N),** Y axes are normalized relative units. Normalized data were spectral area-based. Some bins had negative Y-axis scale after normalization (Metaboanalyst software analysis). Bar charts represent normalized data (mean +/- one standard deviation). Error bars represent the 5% and 95% percentiles, whereas circles represent single data points. Horizontal lines represent box medians. n=6 to 7 biological replicates per group.

### Figure S3

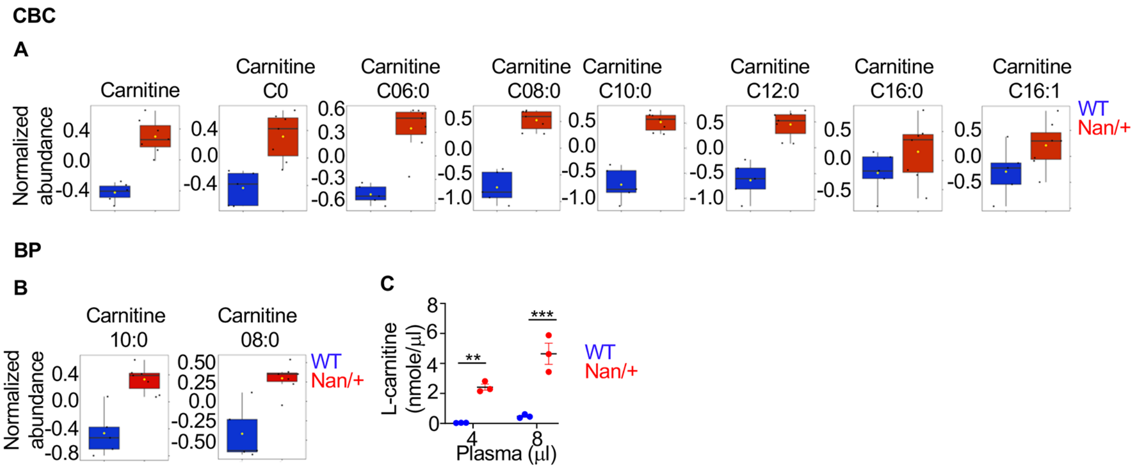


**Figure S3. Carnitine levels in adult circulatory blood cells (CBC) and plasma (PB), in Nan/+ adult mice**

**(A-B)** Box plots of normalized abundance concentration of L-carnitine levels significantly different in WT (blue bar) and Nan/+ (red bar) **CBC** ($p < 0.05$; range = 0.031078 to 4.66E-06) **(A)** and **BP** ($p < 0.05$; range = 0.042103 to 0.00043188) **(B)** obtained from WT or Nan/+. Y axes are normalized relative units. Normalized data were spectral area-based. Some bins had negative Y-axis scale after normalization (Metaboanalyst software analysis). Bar charts represent normalized data (mean +/- one standard deviation). Error bars represent the 5% and 95% percentiles, whereas circles represent single data points. Horizontal lines represent box medians. n=5 biological replicates per group.

**(C)** Concentration of L-carnitine levels in **BP** of WT or Nan/+ adult mouse were analyzed by an ELISA assay. Data are expressed as the mean ± S.E.M. from n=4 biological replicates per group, **$P < 0.01$, ***$P < 0.001$.

### Figure S4

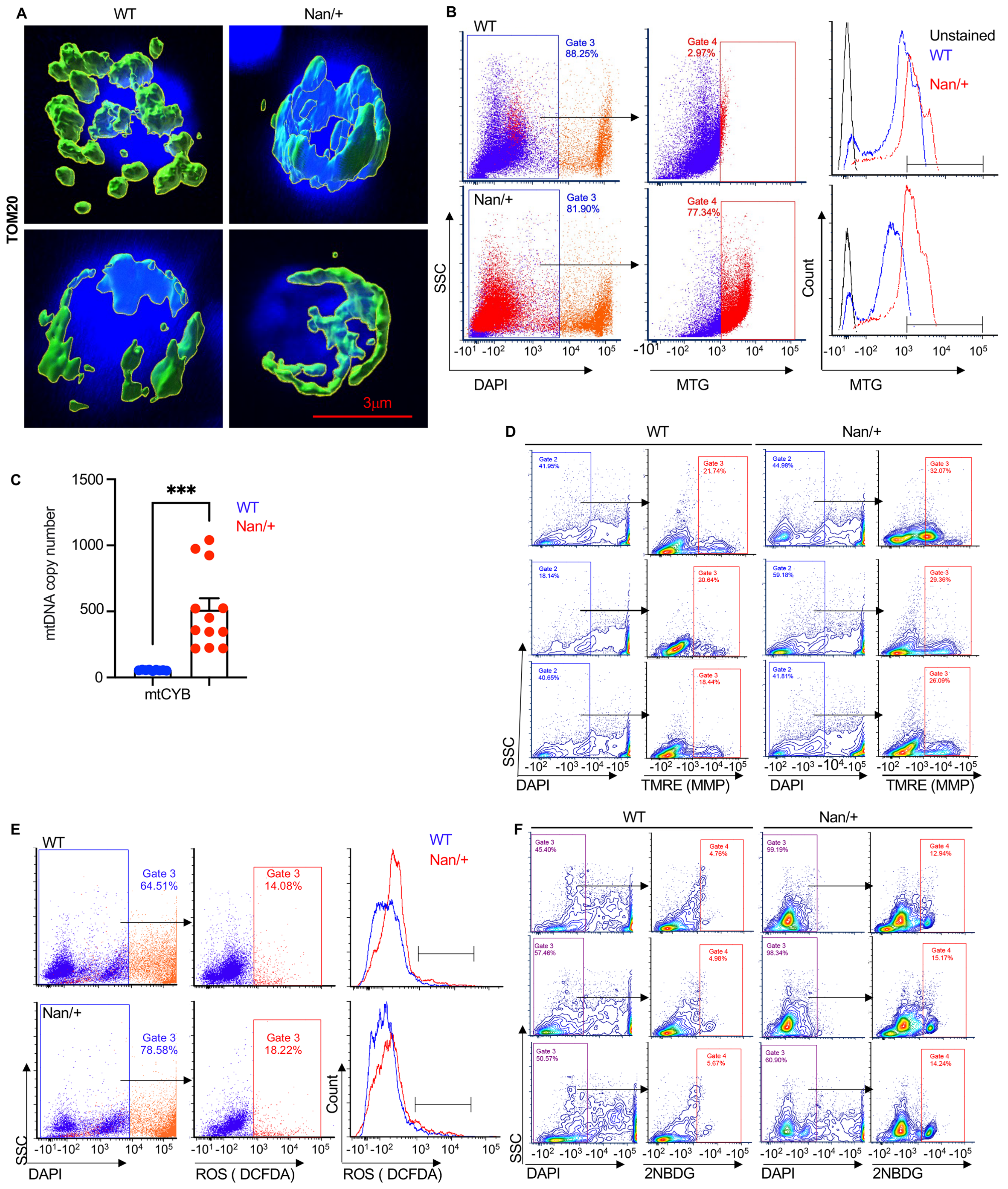
A
WT
Nan/+
TOM20
3μm
B
WT
Gate 3
88.25%
Gate 4
2.97%
Unstained
WT
Nan/+
Nan/+
Gate 3
81.90%
Gate 4
77.34%
SSC
Count
DAPI
MTG
MTG
C
***
WT
Nan/+
mtDNA copy number
1500
1000
500
0
mtCYB
D
WT
Nan/+
SSC
DAPI
TMRE (MMP)
DAPI
TMRE (MMP)
E
WT
Gate 3
64.51%
Gate 3
14.08%
WT
Nan/+
Nan/+
Gate 3
78.58%
Gate 3
18.22%
SSC
Count
DAPI
ROS ( DCFDA)
ROS ( DCFDA)
F
WT
Nan/+
SSC
DAPI
2NBDG
DAPI
2NBDG

**Figure S4.** Mitochondrial morphology and properties are altered in Nan/+ FL cells

**(A)** Representative immunofluorescent confocal images of mitochondrial protein TOM20 showing the morphology of mitochondria (bar= 3 μm) (Related to Fig. 2A).

**(B)** Gating strategy used for analyzing mitochondrial mass in freshly isolated FL cells from WT or Nan/+ measured by MitoTracker Green (left), and representative histogram of mitochondrial mass (right) (Related to Fig. 2B).

**(C)** Mitochondrial DNA copy numbers (mtDNA) were determined using RT-qPCR-based analyses to measure the ratio of mitochondrially encoded Cytochrome B (MT-CYB) versus nuclear ß-actin. Data are presented as mean ± S.E.M. (∗∗∗p < 0.001, (Related to Figure 2C).

**(D)** Gating strategy (n=4 mice) used to identify mitochondrial membrane potential levels (MMP) and representative flow contour plots of MMP in freshly isolated FL cells from WT or Nan/+ (Related to Fig. 2E).

**(E)** Gating strategy used to identify ROS levels (left), and representative histogram showing ROS levels in freshly isolated FL cells from WT or Nan/+ measured by DCFDA (right; Related to figure 2G).

**(F)** Gating strategy used for analyzing glucose uptake in freshly isolated FL cells from WT or Nan/+ measured by 2NBDG (Related to Fig. 2H). Data are presented from n =3 to 6 biological replicates per group.

### Figure S5

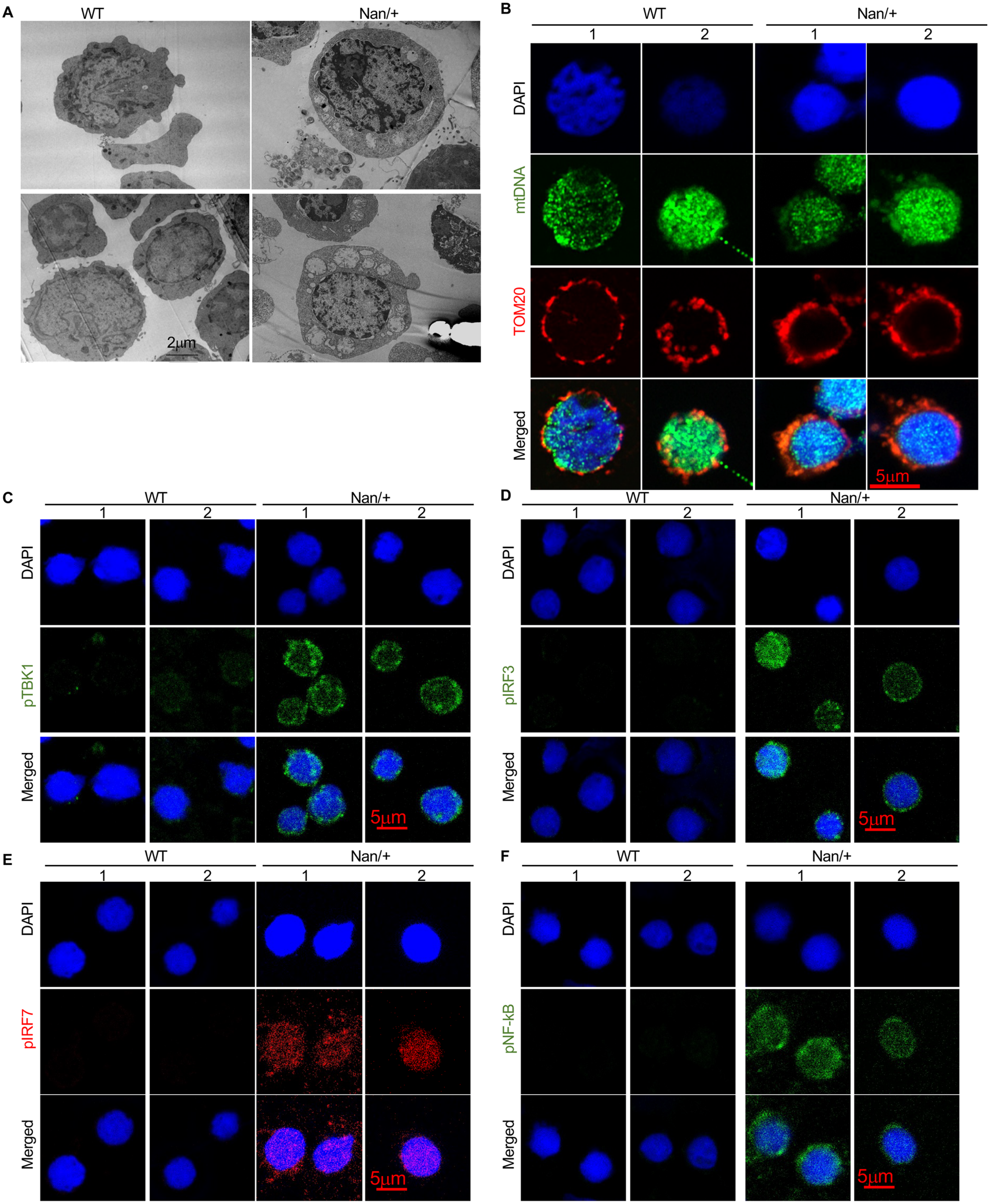

A
WT
Nan/+
2μm
B
WT
Nan/+
1
2
1
2
DAPI
mtDNA
TOM20
Merged
5μm
C
WT
Nan/+
1
2
1
2
DAPI
pTBK1
Merged
5μm
D
WT
Nan/+
1
2
1
2
DAPI
pIRF3
Merged
5μm
E
WT
Nan/+
1
2
1
2
DAPI
pIRF7
Merged
5μm
F
WT
Nan/+
1
2
1
2
DAPI
pNF-kB
Merged
5μm

**Figure S5. STING signaling pathway is activated in Nan/+ FL cells**

**(A)** Representative transmission electron microscopy images of mitochondria in FL cells from WT or Nan/+ FL cells (bar = 2μm) (Related to figure 3A).

**(B)** Representative immunofluorescent super resolution confocal images of mtDNA colocalization with mitochondrial protein TOM20 in freshly isolated FL cells from WT or Nan/+ (bar= 5μm) (Related to Figure 3B).

(**C-F)** Representative confocal images showing immunofluorescent of cGAS-STING pathway related proteins: pTBK1 **(C),** nuclear translocation of pIRF3 **(D),** pIRF7 **(E),** and pNF-kB p65 **(F)** in freshly isolated FL cells from WT or Nan/+ (bar = 5μm) (Related to Fig. 3 G-J). Data are presented from n =3 to 4 biological replicates per group.

### Figure S6

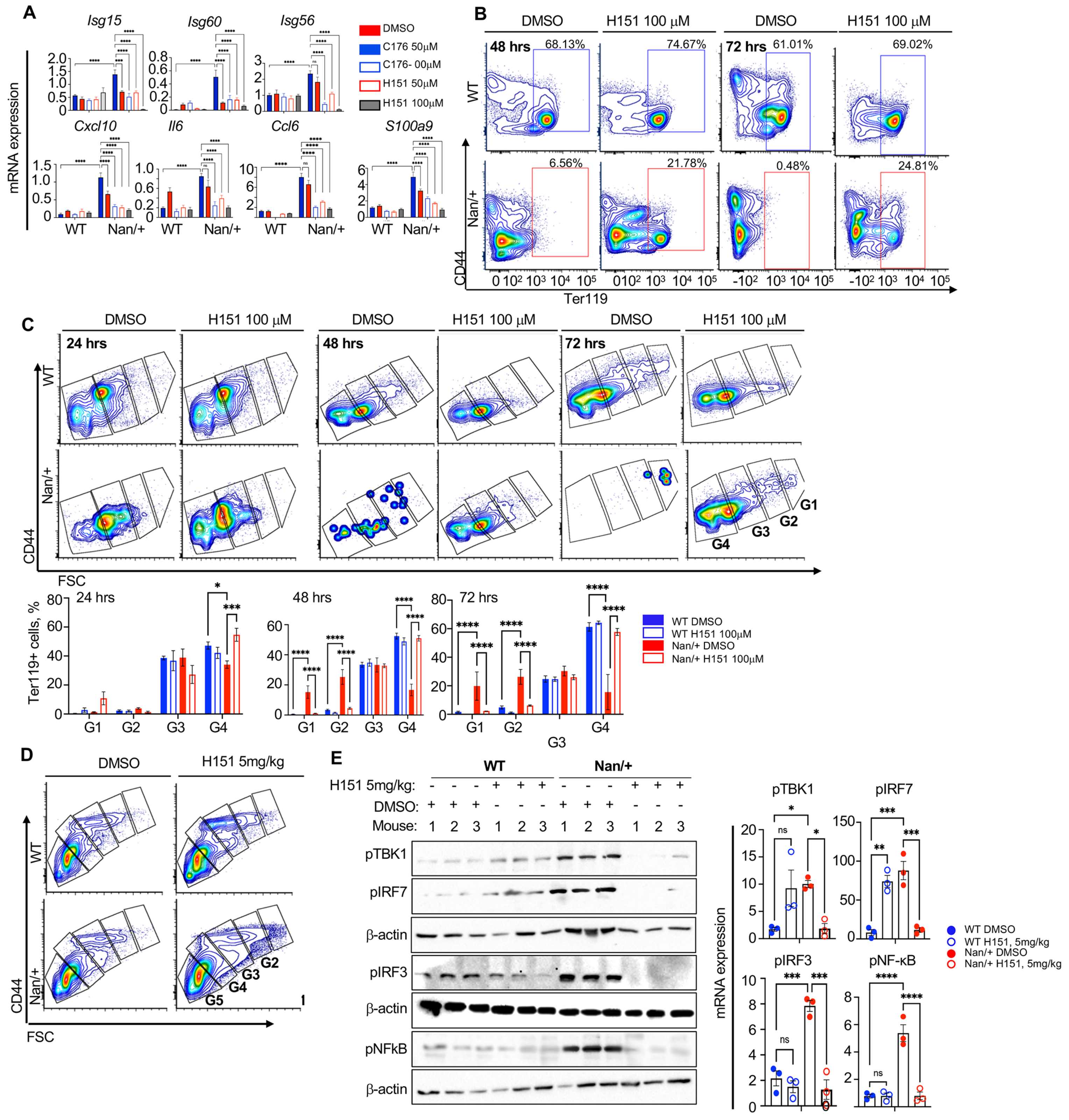
A
Isg15
Isg60
Isg56
Cxcl10
Il6
Ccl6
S100a9
mRNA expression
DMSO
C176 50μM
C176- 00μM
H151 50μM
H151 100μM
WT
Nan/+
B
DMSO
H151 100 μM
48 hrs
72 hrs
68.13%
74.67%
61.01%
69.02%
6.56%
21.78%
0.48%
24.81%
CD44
Ter119
C
24 hrs
FSC
G1
G2
G3
G4
Ter119+ cells, %
WT DMSO
WT H151 100μM
Nan/+ DMSO
Nan/+ H151 100μM
D
H151 5mg/kg
G5
E
H151 5mg/kg:
DMSO:
Mouse:
pTBK1
pIRF7
β-actin
pIRF3
pNFkB
pNF-κB
WT H151, 5mg/kg
Nan/+ H151, 5mg/kg

**Figure S6: STING inhibitor ameliorates anemia in Nan/+**

**(A)** Relative mRNA levels of ISGs and inflammation related genes from WT or Nan/+ FL cells (normalized to β-actin) treated with STING inhibitors (normalized to β-actin) (Related to Fig.4A, bottom).

(**B**) Flow plots showing the gating strategy used to identify CD44/Ter119 cells, in freshly isolated FL cells from WT or Nan/+ and treated in time dependent manner (24- 72hrs) with STING inhibiter (H151, 100 μM) (Related to Fig. 4B).

**(C)** Representative flow plots (top) showing the gating strategy used for analysis of erythroid populations in FL cells treated with DMSO or STING inhibitor (H151, 100 μM) in a time dependent manner (24- 72hrs) , and gated for G1-G4 as indicated (bottom).

**(D)** Representative flow plot showing the gating strategy used for analysis of bone marrow cells isolated from DMSO or STING inhibitor (H151, 5mg/kg) treated mice and were analyzed for red cell maturation (Related to Fig. 4D)

**(E)** Western blot analysis of the indicated proteins that play a role in mDNA-cGAS- STING was performed using isolated Ter119 + cells from the BM of DMSO or Nan/+ adult mice treated with STING inhibitor (H151, 5mg/kg) (**Left,** n=3). Their quantification is shown in the **right**. β-actin was used as a loading/normalization control.

In all panels, data are presented as mean ± S.E.M. (*$p < 0.05$, **$p < 0.01$, and ***$p < 0.001$, and ****$p < 0.0001$).

### Figure S7

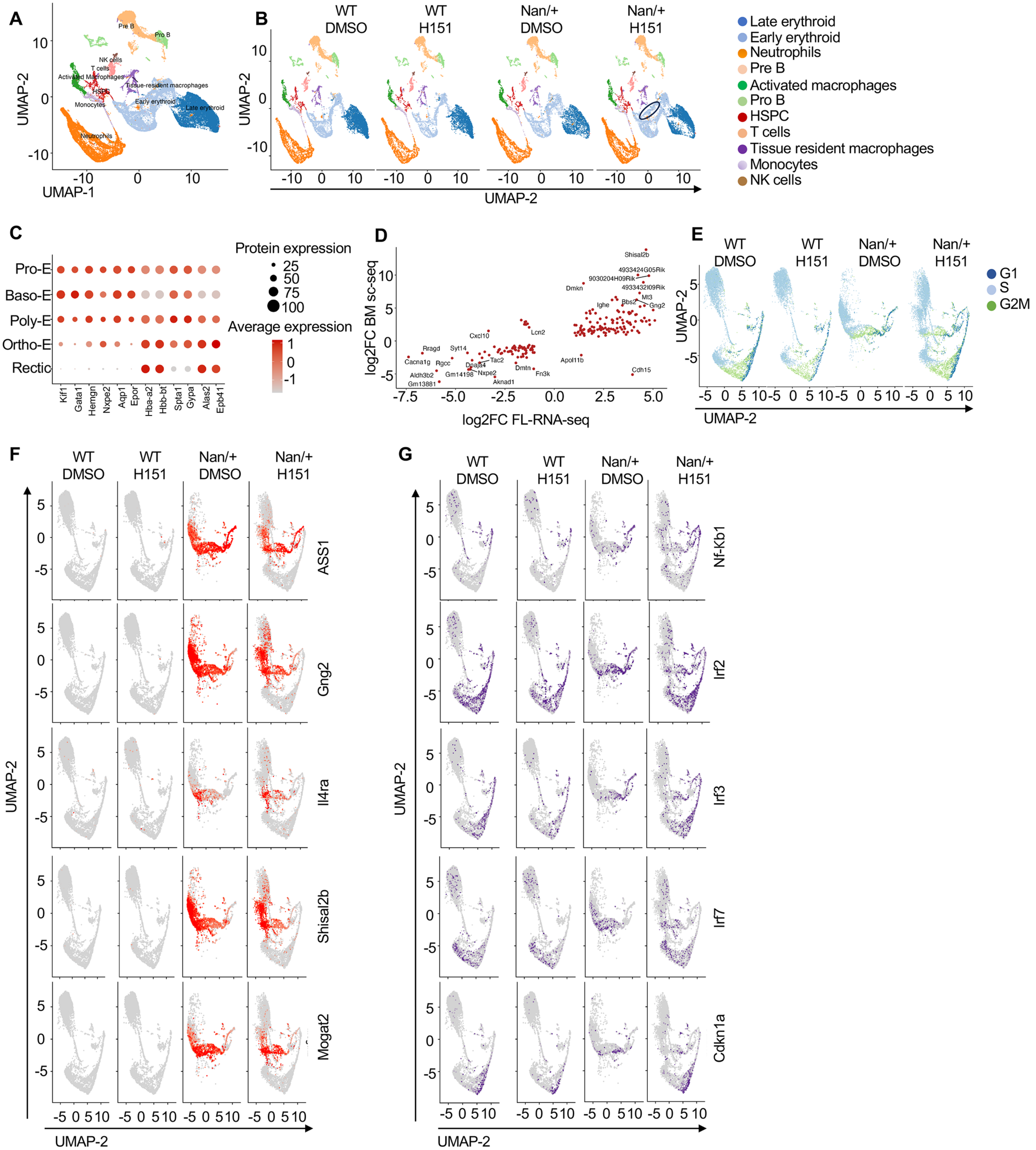
A
UMAP-2
UMAP-1
Pre B
Pro B
NK cells
T cells
Activated Macrophages
HSPC
Tissue-resident macrophages
Monocytes
Early erythroid
Late erythroid
Neutrophils
B
WT DMSO
WT H151
Nan/+ DMSO
Nan/+ H151
UMAP-2
Late erythroid
Early erythroid
Neutrophils
Pre B
Activated macrophages
Pro B
HSPC
T cells
Tissue resident macrophages
Monocytes
NK cells
C
Pro-E
Baso-E
Poly-E
Ortho-E
Rectic
Kif1
Gata1
Hemgn
Nxpe2
Aqp1
Epor
Hba-a2
Hbb-bt
Spta1
Gypa
Alas2
Epb41
Protein expression
25
50
75
100
Average expression
D
log2FC BM sc-seq
log2FC FL-RNA-seq
Shisal2b
4933424G05Rik
9030204H09Rik
4933432I09Rik
Dmkn
Mt3
Bbs2
Gng2
Ighe
Lcn2
Cxcl10
Rragd
Syt14
Cacna1g
Rgcc
Tac2
Dnaja4
Aldh3b2
Gm14198
Nxpe2
Gm13881
Aknad1
Dmtn
Fn3k
Apol11b
Cdh15
E
WT DMSO
WT H151
Nan/+ DMSO
Nan/+ H151
G1
S
G2M
UMAP-2
F
WT DMSO
WT H151
Nan/+ DMSO
Nan/+ H151
ASS1
Gng2
Il4ra
Shisal2b
Mogat2
UMAP-2
G
WT DMSO
WT H151
Nan/+ DMSO
Nan/+ H151
Nf-Kb1
Irf2
Irf3
Irf7
Cdkn1a
UMAP-2

**Figure S7. Single cell analysis of WT and Nan/+ bone marrow erythroid population.**

**(A)** Clustering and U-MAP projection of single cell RNA-Seq data from pooled BM cells from WT and Nan/+ mice with and without H151 treatment. Clusters are colored by hematopoietic cell-type and labeled accordingly.

**(B)** Clustering and U-MAP projection of bone marrow populations of samples separated by genotype with and without H151 treatment. Black oval shows a "new" erythroid population in Nan/+ samples treated with H151.

**(C)** Dot plot showing expression of erythroid markers in the clusters identified in (Fig. 1A).

**(D)** Scatterplot showing the log fold change comparison of differentially expressed genes in WT and Nan/+ E13.5 fetal liver from ref {Planutis, 2017 #2297} and WT and Nan/+ erythroid adult bone marrow cells expressing EKLF (this study).

**(E)** Cell cycle scoring and U-MAP projection of erythroid populations separated by sample and labeled based on the cell cycle stage.

**(F)** Feature plot showing the expression of specific ectopically expressed genes.

**(G)** Feature plot showing the expression of STING target genes involved in Type I inflammatory response.

### Figure S8

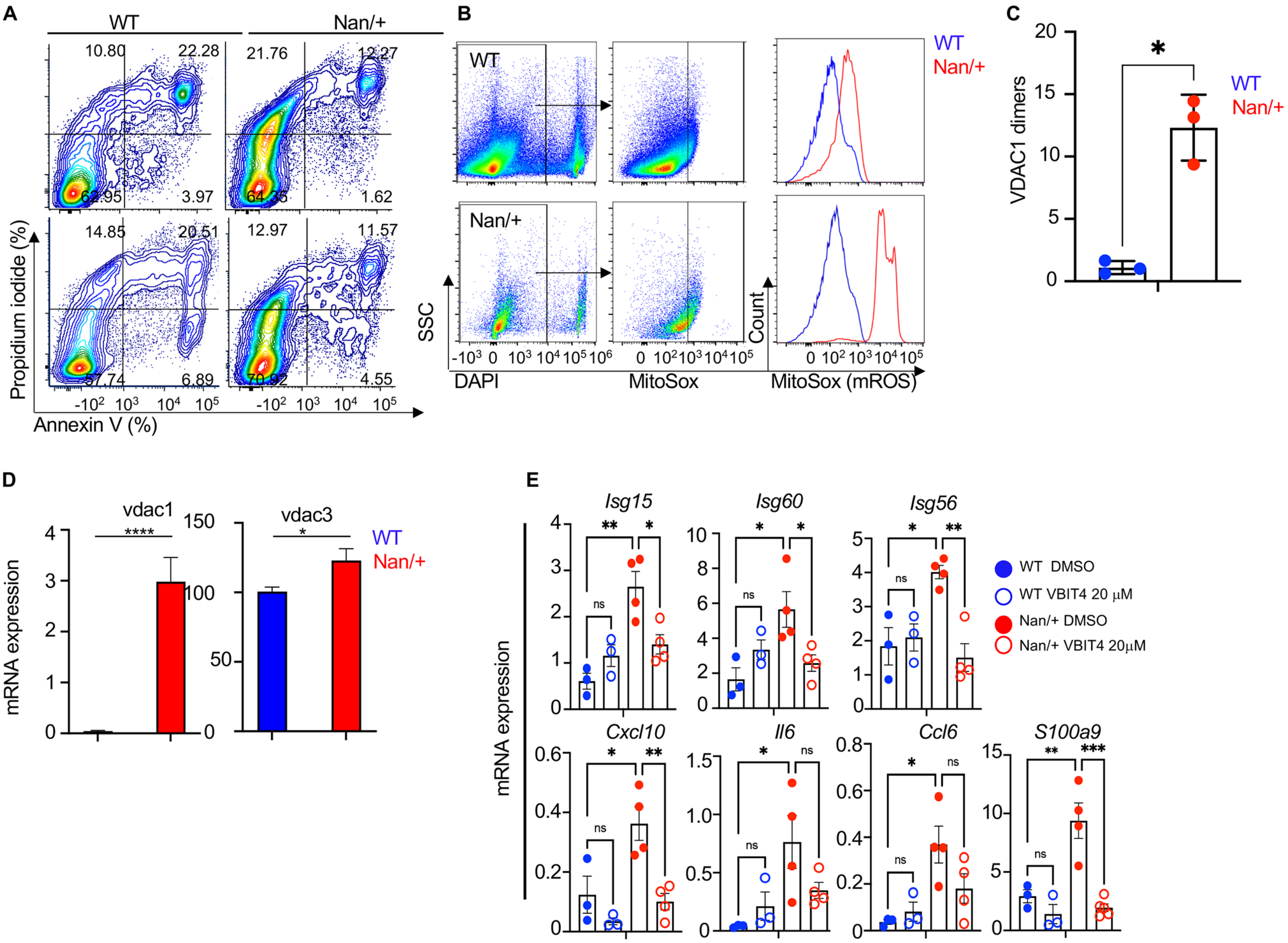


**Figure S8. Mitochondria stress contributes to inflammation in Nan/+ FL cells**

**(A)** Representative flow plot showing apoptotic cell death analyzed by staining with annexin-V FITC/propidium iodide (related to Fig. 7A).

**(B)** Gating strategy used for analyzing mitochondrial ROS (mROS) in freshly isolated FL cells from WT or Nan/+ measured by Mitosox (left), and representative histogram (right) (related to Fig. 7B).

(**C**) Quantification of VDAC1 oligomerization levels were shown without EGS-base cross-linking and probing with anti-VDAC1 antibody. VDAC1 monomers, dimers, and multimers are as indicated in Fig. 7C. and VDAC1 dimer levels were quantified as relative units compared to WT (related to Fig 7C).

**(D)** RT-qPCR analysis of VDAC1 and VDAC3 transcripts in FL cells presented in FL cells from WT, or Nan/+ (normalized to ß-actin). Data are presented as mean ± S.E.M. from n =3 to 4 biological replicates per group (*p < 0.05 and ****p < 0.0001).

(**E**) Relative mRNA levels of ISGs and inflammation related genes from WT and Nan/+ FL cells treated with VBIT4 (20 μM) for 24 hrs (normalized to β-actin) are as indicated (related to Fig.7G).

In all panels, data are presented as mean ± S.E.M. (*p < 0.05, **p < 0.01, and ***p < 0.001, and ****p < 0.0001
